# Reward enhances memory via age-varying online and offline neural mechanisms across development

**DOI:** 10.1101/2021.09.14.460286

**Authors:** Alexandra O. Cohen, Morgan M. Glover, Xinxu Shen, Camille V. Phaneuf, Kristen N. Avallone, Lila Davachi, Catherine A. Hartley

**Affiliations:** Department of Psychology, New York University, New York, NY 10003; Department of Psychology, Columbia University, New York, NY 10027; Nathan Kline Institute of Psychiatric Research, Orangeburg, NY 20962; New York University Center for Neural Science and Langone Health Neuroscience Institute, New York, NY 10003

## Abstract

Reward motivation enhances memory through interactions between mesolimbic, hippocampal, and cortical systems — both during and after encoding. Developmental changes in these distributed neural circuits may lead to age-related differences in reward-motivated memory and the underlying neural mechanisms. Converging evidence from cross-species studies suggests that subcortical dopamine signaling is increased during adolescence, which may lead to stronger memory representations of rewarding, relative to mundane, events and changes in the contributions of underlying subcortical and cortical brain mechanisms across age. Here, we used fMRI to examine how reward motivation influences the “online” encoding and “offline” post-encoding brain mechanisms that support long-term associative memory from childhood to adulthood in human participants of both sexes. We found that reward motivation led to both age-invariant enhancements and nonlinear age-related differences in associative memory after 24 hours. Furthermore, reward-related memory benefits were linked to age-varying neural mechanisms. During encoding, interactions between the prefrontal cortex and ventral tegmental area (VTA) were associated with better high-reward memory to a greater degree with increasing age. Pre- to post-encoding changes in functional connectivity between the anterior hippocampus and VTA were also associated with better high-reward memory, but more so at younger ages. Our findings suggest that there may be developmental differences in the contributions of offline subcortical and online cortical brain mechanisms supporting reward-motivated memory.

**Significance Statement:** A substantial body of research has examined the neural mechanisms through which reward influences memory formation in adults. However, despite extensive evidence that both reward processing and associative memory undergo dynamic change across development, few studies have examined age-related changes in these processes. We found both age-invariant and nonlinear age-related differences in reward-motivated memory. Moreover, our findings point to developmental differences in the processes through which reward modulates the prioritization of information in long-term memory – with greater early reliance on offline subcortical consolidation mechanisms and increased contribution of systems-level online encoding circuitry with increasing age. These results highlight dynamic developmental changes in the cognitive and neural mechanisms through which motivationally salient information is prioritized in memory from childhood to adulthood.

## Introduction

Memories of rewarding experiences can adaptively facilitate the pursuit of rewards across the lifespan. Cross-species research has demonstrated that reward motivation enhances memory in adults (Shohamy and Adcock, 2010; Murty and Dickerson, 2016). Although recent work suggests that children’s memory is also sensitive to value associations during encoding (Ngo et al., 2019), both reward processing and associative memory undergo dynamic changes across development (Ghetti and Bunge, 2012; Galván, 2013; Meyer and Pattwell, 2020). Because memories guide thoughts and actions, insights from studies examining how reward motivation influences memory formation across age can be leveraged to promote healthy development. While considerable research has examined how rewards influence decision-making across adolescence (Galván, 2013; Steinberg et al., 2015; Davidow et al., 2018), few studies have examined how reward influences memory from childhood to adulthood.

Central to reward-motivated memory mechanisms are mesolimbic dopaminergic pathways that originate in the ventral tegmental area (VTA) and project diffusely throughout the brain (Haber and Knutson, 2010; Shohamy and Adcock, 2010). During encoding, top-down modulation of the mesolimbic system by the prefrontal cortex (PFC) facilitates reward-motivated memory (Miendlarzewska et al., 2016; Murty and Dickerson, 2016). Interactions between lateral PFC and VTA have been implicated in driving reward-motivated behaviors and memory formation (Ballard et al., 2011; Murty and Adcock, 2014). Increased VTA and anterior hippocampus activation and functional connectivity during encoding have also been linked to reward-motivated memory (Adcock et al., 2006).

Beyond “online” encoding, “offline” post-encoding consolidation mechanisms stabilize information in long-term memory (Tambini et al., 2010; Tambini and Davachi, 2019). Encoding and post-encoding processes make dissociable contributions to reward-motivated associative memory (Murty et al., 2017). Moreover, reward-related memory enhancements typically emerge after a delay, underscoring the importance of offline post-encoding mechanisms for prioritizing high-reward associations in memory (Miendlarzewska et al., 2016; Dickerson and Adcock, 2018). Changes in VTA and anterior hippocampus functional connectivity following reward-motivated encoding have been linked to better high-reward associative memory (Gruber et al., 2016; Murty et al., 2017). In sum, reward-motivated memory formation is supported by complementary online and offline mechanisms comprising distributed neural circuits.

Developmental changes in mesolimbic system function may yield corresponding changes in reward-related memory processes. Cross-species evidence (Haycock et al., 2003; Weickert et al., 2007s; Tseng and O’Donnell, 2007; Brenhouse et al., 2008) suggests that subcortical dopamine signaling increases during adolescence. Such developmental changes in mesolimbic system function have been linked to adolescents’ increased risky, impulsive, and effortful behaviors in response to rewards (Galván, 2013; Luna et al., 2015; Doremus-Fitzwater and Spear, 2016). Given the central role of dopamine-dependent plasticity and VTA functional connectivity in adult memory formation (Huang and Kandel, 1995; Li et al., 2003; Morris et al., 2003; Duncan et al., 2014; Tompary et al., 2015), nonlinear developmental changes in mesolimbic system function may lead to more robust reward-related memory during adolescence.

Dynamic developmental changes in brain circuitry may also lead to age-related shifts in reward-motivated memory mechanisms. Theories of brain development posit that subcortical circuitry functionally matures in childhood, whereas broader circuits that include PFC show protracted development into adulthood (Casey et al., 2016, 2019). One study in adolescents and adults indicated that same-day positive memory biases in adolescents are related to enhanced hippocampal-striatal functional connectivity during learning (Davidow et al., 2016). These findings suggest that reliance on subcortical reward memory mechanisms may be greater earlier in development. Moreover, offline consolidation during sleep is particularly beneficial for learning and memory in children (Kurdziel et al., 2013, 2018; Wilhelm et al., 2013). Still, the contributions of encoding and post-encoding brain activity to reward-motivated memory across development have not been studied.

Here, we examine how reward associations influence long-term associative memory from childhood to adulthood and whether there are age-related differences in the underlying online and offline neural mechanisms. Given prior work suggesting developmental changes in memory specificity (Keresztes et al., 2018), we examined both general and specific associative memory measures and their associations with encoding- and post-encoding-related brain activation and functional connectivity across age. We hypothesized that reward motivation may uniquely facilitate associative memory during adolescence due to enhanced post-encoding subcortical functional connectivity and that top-down prefrontal encoding mechanisms may strengthen with age.

## Materials and Methods

### Participants

Eighty-nine participants ages 8- to 25-years-old (M_age_ = 16.16, SD_age_ = 4.67, 45 female) were included in analyses. A target sample size of *n* = 90, including 30 children, 30 adolescents, and 30 adults, was determined based on prior work using similar or smaller sample sizes to identify age-related differences in behavior and brain activation (Van Den Bos and Rodriguez, 2015; Insel et al., 2019; Callaghan et al., 2021). Data exclusions consisted of: eight participants with excessive motion (participants without at least one complete encoding, baseline arrows, and post-encoding arrows runs due to exclusions of runs with 15% or more timepoints censored with greater than 0.9 mm relative translational motion), seven participants who elected to not complete or terminate the fMRI scan, and five participants with incomplete datasets as a result of fMRI scanner malfunction. Participants comprised a sample of volunteers recruited from the local community of New York City. Of the 89 participants included in analyses, 12.4% identified as African American/Black, 24.7% as Asian, 38.2% as Caucasian/White, 1.1% as Native American, and 23.6% as more than one race. In addition, 15.7% of the sample identified as Hispanic. Based on self or parental report, participants were right-handed and reported no: previous head injury, serious neurological or medical illness, diagnosed psychiatric illness, developmental disability, sensory impairment such as vision or hearing loss, use of medications that influence the functioning of the central nervous system or peripheral physiological responses (e.g., beta-blockers), or major contraindication for MRI. Participants ages 18 and over provided informed written consent and minor participants provided assent, according to research procedures approved by New York University’s Institutional Review Board. Parents or guardians of participants under age 18 also provided written consent on behalf of the child prior to participation in the study. Participants were compensated $75 and up to $21 in bonus money for their participation in two sessions.

### Experimental design and statistical analyses

Participants completed a high- and low-reward-motivated encoding and retrieval task, adapted from previous work (Duncan et al., 2014; Murty et al., 2017), with baseline and post-encoding active rest periods in the fMRI scanner (Figure 1). Approximately one week prior to the scan, child and adolescent participants completed a mock scan to acclimate to the scanning environment and to practice remaining as still as possible while inside the mock scanner. Immediately prior to the scan, participants received instructions about each component of the scan and completed a practice session including four encoding and four retrieval trials using images that were not included in the fMRI tasks. Tasks were programmed in Expyriment (Krause and Lindemann, 2014) using Pygame v1.9.4 and Python v2.7.15. The reward-motivated encoding task included images from RADIATE (Conley et al., 2018), the Chicago Face Database (Ma et al., 2015), Harvard’s Konkle Lab (Konkle et al., 2010), and MIT’s Places Scene Recognition (Zhou et al., 2014) databases. Following the practice session, participants completed a three-minute “pre-exposure” to the eight source images (four faces and four places, repeated five times each and for three-second presentations) that would be included in the motivated encoding task in order to familiarize participants with these repeating images and mitigate any potential effects of source image category on memory performance (Mayes et al., 2007). Participants returned 24 hours after the first session to complete a behavioral retrieval task.

**Figure 1.**
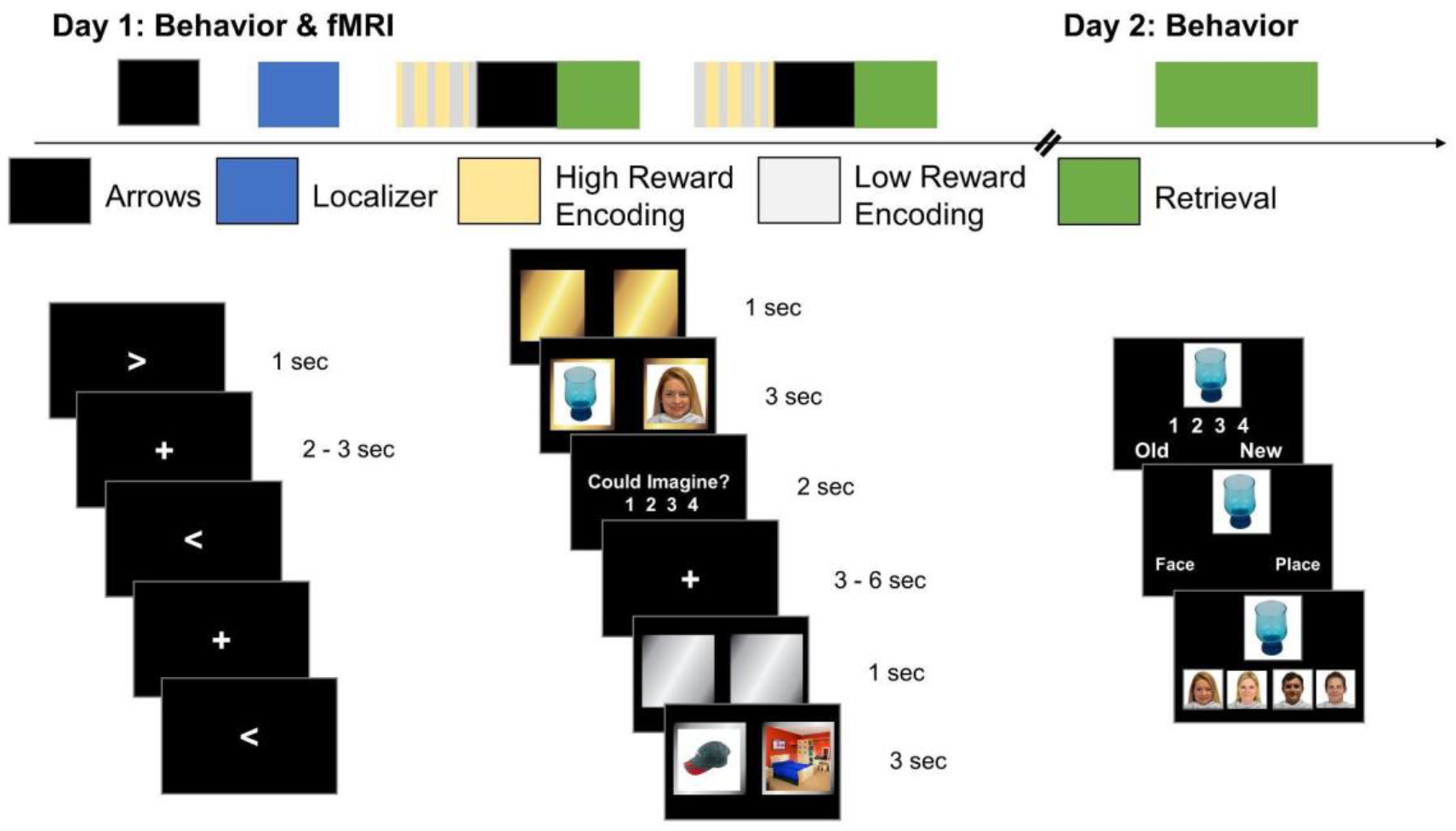
Experimental Design. On day one, participants completed a high and low reward-motivated encoding fMRI task. Participants also completed a baseline and post-encoding arrow detection task (active rest). Participants returned to the lab 24 hours later to complete a memory retrieval task including all the paired associates from day one. For visualization of the day one retrieval phase that was not analyzed for the purposes of this study, see Figure 1-1 in Extended Data.

### Arrows (active rest) task

Participants first completed a 5.57-minute baseline active rest or arrows scan. In the arrows task, participants were instructed to press one button if the arrow pointed to the left (<), or another button if the arrow pointed to the right (>). An “active” rest with a mildly engaging, low cognitive load task was used due to previous work suggesting that active rests serve as better rest measurements for memory studies as compared to passive rest (Stark and Squire, 2003). The arrows task consisted of 94 randomized trials (one second each) with a randomized, jittered intertrial interval (ITI) of two to three seconds. Participants completed a baseline arrows run and two additional arrows runs, following the first and second rounds of encoding.

### Reward-motivated encoding task

The first motivated encoding scan followed a localizer (not analyzed here). On each trial, participants first saw two squares that were either gold or silver, indicating that remembering the upcoming pair of images would help them win a big bonus of $15 (gold high-reward) or a small bonus of $1 (silver low-reward). After one second, a trial-unique picture of an object (selected to be equivalently familiar to participants of all ages) was overlaid on the left square and one of eight repeated source images was overlaid on the right square for three seconds. Source images consisted of four faces (two women, two men) and four scenes (two indoor, two outdoor). Participants were assigned faces or places as the high-reward category of images based on participant ID number. Participants were instructed to create a story involving both images to promote deep encoding. To help maintain attention, participants had two seconds to rate how well they imagined the story on a scale from one (very easy to imagine) to four (very hard to imagine; see Extended Data Figure 2-2 for imagine ratings). Each six second trial was followed by a randomized, jittered ITI of three-six seconds, determined based on previous studies (Mumford, 2014; Mumford et al., 2014). Each encoding phase consisted of 64 trials (32 high- and 32 low-reward) and lasted 11.37 minutes. After encoding, participants completed a post-encoding arrows task, as described above. Finally, participants completed a retrieval scan including half of the trial-unique objects from the preceding encoding block which lasted 6.57 minutes. Day one retrieval data were not analyzed for the purposes of this study (see Extended Data Figure 1-1 for visualization of the day one retrieval phase). Participants repeated another round of encoding, arrows, and retrieval.

### Memory retrieval test

Participants completed a memory retrieval test 24 hours later. We queried retrieval one day later based on prior work suggesting that emotional memory enhancement effects typically emerge with time. Participants were presented with all 128 of the objects observed during encoding phases of the scanning task. Half of these images had also been viewed during the retrieval phases of the scanning task and half of the images were only viewed once during encoding. Additionally, 128 new objects were included as lure images (256 trials total). On each trial, participants saw one object. They first indicated if the object was definitely old, maybe old, maybe new, or definitely new. If they endorsed the object as “definitely old” or “maybe old” they then indicated whether that object was paired with a face or scene. Participants then selected a specific source image. If the participant endorsed the object as “new”, they were not asked further questions about that object. The retrieval test was self-paced.

### Behavioral data analyses

Behavioral data processing and statistical analyses were conducted using R version 3.5.1 (Team, 2016). Mixed-effects models were run using the “lme4” package (version 1.1-17) (Bates et al., 2011). Age was treated as a continuous variable in all analyses and was z-scored across all participants. We examined three components of long-term associative memory — general, specific, and gist source memory. General source hits were defined as the trials where the correct category of source image (i.e., face or place) was identified. Specific source hits were defined as the trials where the correct specific source image (i.e., the specific woman, man, indoor place, or outdoor place) was identified. Gist source hits were defined as trials where the correct category of source image was identified but did not include trials where the correct specific source image was identified (i.e., the difference between general and specific source hits). Because associative memory was only queried on items identified as old, the denominator for these measures was computed as the total number of items correctly identified as old for each given participant. Thus, all of the associative memory measures have the same denominator within each participant. Trials were subdivided into paired associates that had been retrieved on both days or only on day two to account for effects of testing on memory performance (see Extended Data Figure 2-1 for visualization of how the memory measures were computed).

For each memory performance measure, we fit models using a mean-centered, scaled linear age predictor and a squared mean-centered, scaled age term to test for nonlinear effects of age. We compared the linear and quadratic models by likelihood ratio chi-square test to select the best fitting models. Specific source memory was not better fit by the quadratic age model (*χ*^2^(4) = 1.98, *p* = 0.74) while general and gist source memory were better fit by quadratic age models (general: *χ*^2^(4) = 11.83, *p* = 0.019; gist: *χ*^2^(4) = 13.19, *p* = 0.010). We fit maximal models, including a single random intercept per participant and random slopes for within-subject fixed effects (reward level [high or low] and retrieval condition [retrieved day 1 or not tested]) and their interaction, and simplified models that failed to converge by removing random slopes until we identified the most complex random effects structure supported by the data (Bates et al., 2015). In addition to a random intercept, all three models included random slopes for retrieval condition. The high reward category of source image (face or place) for each subject was also included as a covariate of no interest in all analyses. Statistical significance of the fixed effects is reported from the analysis of variance (Type III using Satterthwaite’s method) performed on lmer models for general, specific, and gist source memory.

### Brain-behavior relation analyses

We examined relations between our neural measures of interest and both high-reward specific and gist source memory benefits due to the different age-related effects observed in these non-overlapping components of associative memory. High-reward specific and gist source memory benefit measures were computed by subtracting low reward memory performance from high reward memory performance. These reward memory benefit measures were used as the dependent variables in separate multiple linear regressions assessing brain-behavior relationships, which included the brain measure of interest and mean-centered, scaled age as predictors. The high reward category of source image (face or place) for each subject was included as a covariate of no interest in all analyses and if a significant main effect was observed, it was treated as a variable of interest.

### MRI data acquisition and preprocessing

Participants were scanned using a 3 Tesla Siemens Prisma scanner with a 64-channel head coil at the NYU Center for Brain Imaging. Anatomical data were acquired with high-resolution, T1-weighted anatomical scans using a magnetization-prepared rapidly acquired gradient echo (MPRAGE) sequence (TR = 2.3s, TE = 2.3ms, TI = .9s; 8° flip angle; .9-mm isotropic voxels, field of view = 192 x 256 x 256 voxels; acceleration: GRAPPA 2 in the phase-encoding direction, with 24 reference lines) and T2-weighted anatomical scans using a 3D turbo spin echo (TSE) sequence (T2: TR = 3.2s, TE = 564ms, Echo Train Length = 314; 120° flip angle, .9-mm isotropic voxels, field of view = 240 x 256 x 256 voxels; acceleration: GRAPPA 2×2 with 32 reference lines in both the phase- and slice-encoding directions). Functional data were acquired with a T2*-weighted, multi-echo EPI sequence with the following parameters: TR=2s, TEs=12.2, 29.48, 46.76, 64.04ms; MB factor = 2; acceleration: GRAPPA 2, with 24 reference lines; effective echo spacing: .245 ms; 44 axial slices; 75° flip angle, 3-mm isotropic voxels, from the University of Minnesota’s Center for Magnetic Resonance Research (Feinberg et al., 2010; Moeller et al., 2010; Xu et al., 2013). Multi-band with multi-echo EPI sequences were used to allow for better denoising of data and reduced signal dropout in subcortical brain regions. Total scan time was approximately 1 hour and 15 minutes, including short breaks between scans.

All anatomical and functional MRI data were preprocessed using fMRIPrep 20.0.6 (Esteban et al., 2019), a robust preprocessing pipeline that adjusts to create the optimal workflow for the input dataset. The default options were used with slice timing disabled and the MNI and T1w output spaces specified. The T1w space functional runs were used as the input file in subsequent analyses. Briefly, anatomical processing steps included intensity nonuniformity correction, skull-stripping, spatial normalization, brain tissue segmentation, and surface reconstruction. FMRIPrep uses the tedana T2* workflow (DuPre et al., n.d.; Kundu et al., 2013) to generate an optimally combined timeseries across all four echoes. This combined timeseries is then used in all subsequent preprocessing steps (e.g., registration estimation of head motion and confounds, susceptibility distortion estimation). All raw and preprocessed data were visually inspected. All subsequent processing and statistical analyses were completed in FSL version 5.0.10 (Jenkinson et al., 2012). Registration matrices were estimated by finding and concatenating the transformations between the T1w functional to structural and structural to MNI space fMRIPrep outputs. Updated registration using these matrices derived from the fMRIPrep outputs were visually inspected. All preprocessed BOLD data were spatially smoothed with a 5-mm FWHM Gaussian kernel in run-level analyses.

### Encoding fMRI analyses

Encoding phase analyses examined encoding-related brain activation for high-relative to low-reward memoranda. Run-level GLMs for each participant had two task regressors: high-reward trials and low-reward trials. Each task regressor was convolved with a double gamma hemodynamic response function and included temporal derivatives. The following nuisance regressors derived from fMRIPrep for each encoding run were also included in these models: six motion (translational and rotational) parameters and their derivatives, a framewise displacement regressor, the first six anatomical noise components (aCompCor), and the cosine components to perform high-pass filtering of the data. Encoding runs were combined via fixed-effects analyses and a group-level mixed-effects analysis was performed using FSL’s FLAME 1. A single group average was calculated for the high reward > low reward contrast-of-interest and included demeaned age and demeaned age-squared as covariates. Whole-brain multiple comparison correction was performed using a *Z*-statistic threshold of *Z* > 3.1 and a cluster-defining threshold of *p* < .05.

A psychophysiological interaction (PPI) analysis was conducted to examine task-dependent connectivity with a dorsolateral prefrontal cortex (dlPFC) region-of-interest (ROI) seed derived from high > low reward contrast (4-mm sphere around peak activation). The functional timecourse within the seed region was extracted from the filtered functional data from the participant-level GLMs. The filtered functional data was also used as input for the participant-level PPI GLMs, which included four regressors: a high-reward - low-reward task regressor, a dlPFC timecourse physiological regressor, a PPI regressor, and a high-reward + low-reward task regressor to model shared variance between the two task conditions. Both task regressors were convolved with a double gamma hemodynamic response function and included temporal derivatives. Connectivity estimates (*Z*-statistics) were extracted for each participant from a VTA ROI (described below). A group-level PPI analysis was also performed as described above.

### ROI definition

We investigated functional connectivity using *a priori* anatomical ROIs that have been previously implicated in reward-motivated memory processes: VTA and anterior hippocampus. We chose to focus on the VTA as dopaminergic midbrain regions have been broadly implicated in memory processes (e.g., Duncan et al., 2014; Tompary et al., 2015) and in particular, reward-motivated memory (e.g., Wittmann et al., 2005; Adcock et al., 2006; Gruber et al., 2016; Murty et al., 2017). The VTA and bilateral anterior hippocampus were defined in standard space using probabilistic atlases that reliably define these areas at conventional MRI resolution (Murty et al., 2014; Ritchey et al., 2015). For post-encoding functional connectivity analyses, these ROIs were warped into subject T1w space and thresholded at 75%. All ROIs were visually inspected.

### Active rest fMRI analysis

Seed-based functional connectivity analyses were performed on active rest data. Run-level GLMs for each participant included a single arrows task regressor convolved with a double gamma hemodynamic response function and including a temporal derivative, all nuisance regressors listed under encoding analyses, and three additional nuisance regressors derived from fMRIPrep for each arrows run: white matter, cerebrospinal fluid, and global signals. The mean residual timecourse for each ROI (VTA and anterior hippocampus) was extracted from each active rest run. Consistent with prior work (Murty et al., 2017), pairwise Pearson correlations were used to examine functional connectivity between the VTA and anterior hippocampus. Correlation scores for each pairwise comparison were averaged across post-encoding scans to obtain a single measure of post-encoding connectivity. Baseline and post-encoding correlation scores were Fisher *r*-to-*z* transformed so that they could be submitted to regressions examining brain-behavior relations (described above). The measures of experience-dependent change in connectivity used in these analyses were obtained by subtracting baseline connectivity from post-encoding.

### Data and code availability

Data & code necessary to reproduce the findings will be made available on Open Science Framework (https://osf.io/7shjq/?view_only=0e9666a83a5546bbbcd093141dfb0ecc) upon publication.

## Results

### Age-invariant and nonlinear components of reward-motivated memory

We examined associative memory measures after 24 hours, given prior work suggesting that reward-motivated memory enhancements in adults typically emerge after a delay (Miendlarzewska et al., 2016; Dickerson and Adcock, 2018) and that associative memory changes with age (Lee et al., 2016, 2020). During reward-motivated encoding on day one (Figure 1), participants encoded paired associates consisting of a trial-unique, child friendly object and either a face or place source image, with each category serving as the high-reward category for half of participants. During retrieval on day two, participants first rated each object as old or new (see Extended Data Figure 2-3 for item recognition memory performance). Then if the object was rated old, participants indicated whether the object had been paired with a face or place, and finally which specific face or place had been paired with the object. We computed general, specific, and gist source memory performance measures from the day two retrieval test. General source hits were defined as the trials where the correct category of source image (i.e., face or place) was identified. Specific source hits were defined as the trials where the correct specific source image (i.e., the specific woman, man, indoor place, or outdoor place) was identified. Gist source hits were defined as trials where the correct category of source image was identified but did not include trials where the correct specific source image was identified (i.e., the difference between general source and specific associative hits; see Materials and Methods and Extended Data Figure 2-1 for additional details).

Using a linear mixed-effects model, we examined more general associative memory performance for whether an object had been associated with a face or place (general source memory) as a function of reward level (high or low), age, age-squared, and retrieval condition (whether the association had been retrieved on both days or only on day two), controlling for the high-reward source image category (whether faces or places were associated with high reward). We fit both linear and quadratic continuous age models to behavioral data and report the results from the best-fitting model for each memory measure (see Materials and Methods). We found significant main effects of reward level (*χ*^2^(1, N = 89) = 7.23, *p* = 0.0072) and a marginally significant effect of age (*χ*^2^(1, N = 89) = 3.11, *p* = 0.078). These main effects were qualified by a significant reward level-by-age-squared interaction (*χ*^2^(1, N = 89) = 10.07, *p* = 0.0015), such that there was a peak in high-reward source memory and a trough in low-reward source memory in adolescence (Figure 2A). There were no significant main effects of age-squared (*χ*^2^(1, N = 89) = 0.74, *p* = 0.39) or high-reward source image category (*χ*^2^(1, N = 89) = 0.13, *p* = 0.71). As expected, we observed a significant effect of retrieval condition (*χ*^2^(1, N = 89) = 5.18, *p* = 0.023) indicating that participants demonstrated better memory for associations that had been retrieved on day one. However, there were no significant interactions of reward level-by-retrieval condition or age, retrieval condition-by-age or age-squared, or reward level-by-retrieval condition by-age or age-squared (all *χ*^2^s < 1.80, all *p*s > 0.18), so we collapsed across retrieval conditions for visualization.

**Figure 2.**
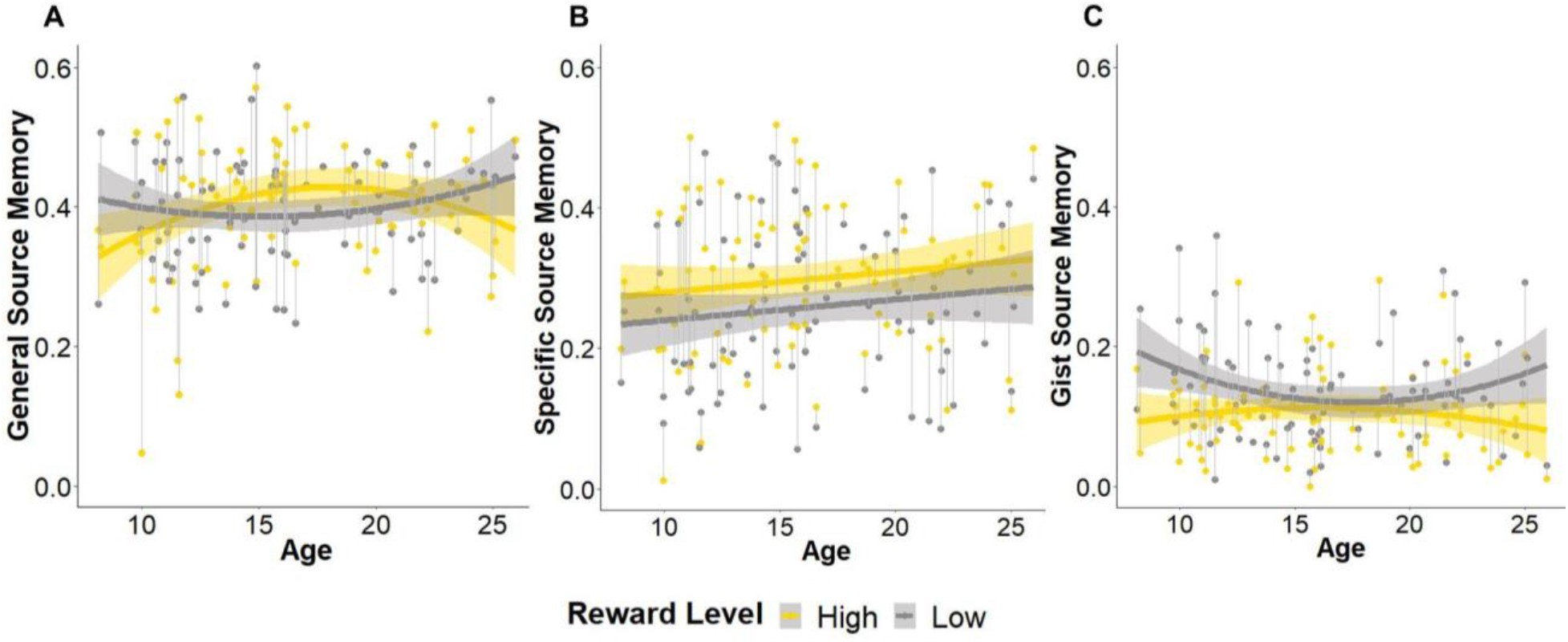
Enhanced specific high-reward associative memory across all ages and nonlinear age-related differences in more general high-reward associative memory during adolescence after 24 hours. (A) General source memory performance shows a nonlinear relationship with age, such that there is a peak in high-reward source memory and a trough in low-reward source memory during adolescence. (B) Across all ages, participants show better memory for specific high-reward relative to low-reward associations. (C) Gist source memory (the difference between general and specific source memory) performance shows a nonlinear relationship with age such that performance is similar for high- and low-reward associations during adolescence but increased for low-reward associations in older and younger participants. Shading depicts a 95% confidence interval around fitted lines. For visualization of how the memory measures were computed, see Figure 2-1 in Extended Data. For encoding phase imagery ratings, see Figure 2-2 in Extended Data. For item recognition memory performance after 24 hours, see Figure 2-3 in Extended Data. For associative memory analyses that include memory confidence level, see Figures 2-4 and 2-5 and Tables 2-1 to 2-3 in Extended Data.

Next, we examined whether participants remembered the specific source image associated with an object (specific source memory) as a function of reward level (high or low), age, and retrieval condition (whether the association had been retrieved on both days or only on day two), controlling for the high-reward source image category (whether faces or places were associated with high reward). We found a significant effect of reward level, such that participants showed better specific source memory for high-reward relative to low-reward memoranda (*χ*^2^(1, N = 89) = 17.89, *p* < 0.001). There was no significant effect of age (*χ*^2^(1, N = 89) = 1.99, *p* = 0.16) or age-by-reward level interaction (*χ*^2^(1, N = 89) = 0.00, *p* = 0.99), suggesting similar reward-motivated memory enhancements across all participants. As expected, we observed a significant effect of retrieval condition (*χ*^2^(1, N = 89) = 49.56, *p* < 0.001) indicating that participants demonstrated better memory for associations that had been retrieved on day one. However, there was no reward level-by-retrieval condition interaction (*χ*^2^(1, N = 89) = 1.85, *p* = 0.17), so we collapsed across retrieval conditions for visualization (Figure 2B). There was no main effect of high-reward source image category and there were no significant interactions of retrieval condition-by-age or reward level-by-retrieval condition by age (all *χ*^2^s < 0.80, all *p*s > 0.35). Thus, individuals showed better memory for specific high-reward relative to low-reward associations across all ages.

Finally, given the overlap between the general and specific source memory measures, we examined correct source memory when the specific source memory was incorrect (gist source memory) using linear mixed-effects analysis as described above. We found a significant reward level-by-age-squared interaction (*χ*^2^(1, N = 89) = 10.21, *p* = 0.0014), such that gist source memory performance was similar for high- and low-reward associations during adolescence but in older and younger participants, low-reward gist source memory performance was increased (Figure 2C). As expected, we observed a significant effect of retrieval condition (*χ*^2^(1, N = 89) = 10.45, *p* = 0.0012), but no reward level-by-retrieval condition interaction (*χ*^2^(1, N = 89) = 0.21, *p* = 0.64), so we again collapsed across retrieval conditions for visualization. There were no other significant main effects or interactions (all *χ*^2^s < 2.60, all *p*s > 0.11).

We also examined how item recognition memory confidence ratings (definitely old or maybe old) influenced associative memory performance and reran each of our associative memory mixed-effects models including confidence level as a fixed-effect. The primary results (i.e., the main effect of reward level in specific source memory and the reward level-by-age-squared interaction effects in general and gist source memory) remain statistically significant (see Extended Data Figures 2-4 and 2-5 and Tables 2-1 to 2-3 for analyses that include memory confidence level).

Taken together, these results showed that reward-motivated associative memory consists of both age invariant and nonlinear age effects.

### Prefrontal encoding activation and functional connectivity with VTA relates to high-reward memory benefits, particularly in older individuals

Whole-brain analysis of encoding-related activation for the high > low reward group contrast revealed increased activation in several areas across the brain including occipital, parietal, and prefrontal cortices as well as the thalamus (see Extended Data Table 3-1; Figure 3A). There were no patterns of activation associated with either age or age-squared covariates. A whole-brain analysis of encoding-related activation for the high > low reward group contrast restricted to trials with correct specific source memory after 24 hours yielded a similar activation map (see Extended Data Figure 3-1, Table 3-2, and Table 3-3). The dorsolateral prefrontal cortex (dlPFC) region from the high > low reward group contrast was of particular interest given previous work indicating protracted development of the prefrontal cortex (Mills et al., 2014) and a role for the dlPFC in driving reward-motivated behaviors (Ballard et al., 2011). We examined the relation between dlPFC activation (mean *Z*-statistic for each participant) and behavioral high-reward memory benefits as a function of age, controlling for the high-reward source image category, using multiple linear regression. We found a significant association between high-reward specific source memory benefits after 24 hours and dlPFC activation across all participants (**β** = 0.035, *t*(84) = 3.01, *p* = 0.0034). There were no significant main effects of age, high-reward source image category, or dlPFC activation-by-age (all *p*s > 0.50). Although regression diagnostics examining influential values did not flag any points as outside of Cook’s distance, the association between dlPFC activation and high-reward specific source memory benefit remains statistically significant when two data points greater than three standard deviations from the mean are excluded from analysis (**β** = 0.028, *t*(82) = 2.05, *p* = 0.044). There were no significant associations between dlPFC activation and high-reward gist source memory benefits after 24 hours (see Extended Data Tables 3-4 & 3-5). These results indicate that increased dlPFC activation was associated with increased high-reward specific source memory benefits after 24-hours across all participants (Figure 3B).

**Figure 3.**
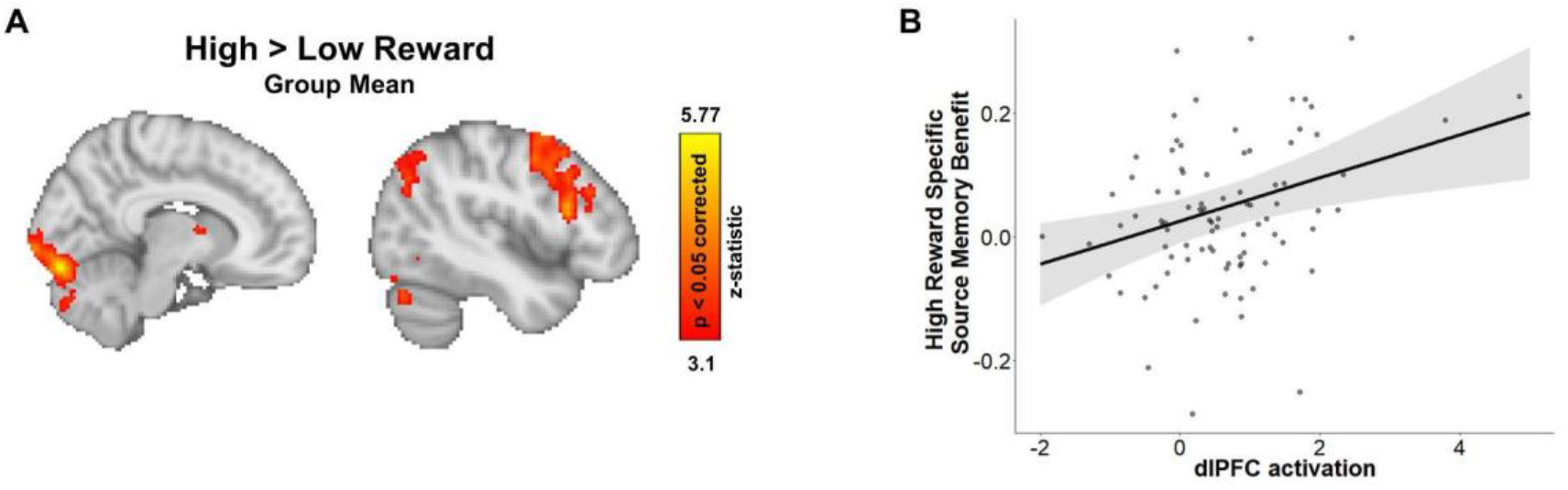
dlPFC encoding activation relates to high-reward memory benefits 24-hours after encoding. (A) Across all participants, brain activation is increased in cortical, thalamic, and cerebellar areas of the brain for encoding high-relative to low-reward stimuli (*p* < 0.05, whole-brain corrected). (B) Increased dorsolateral prefrontal cortex (dlPFC) activation is associated with greater high-reward specific source memory benefits across all participants. Shading depicts a 95% confidence interval around the predicted probability line. For a description of all activations in the high > low reward group contrast, see Table 3-1 in Extended Data. For results of the whole-brain analysis of encoding-related activation for the high > low reward group contrast restricted to trials with correct specific source memory after 24 hours, see Figure 3-1, Table 3-2, and Table 3-3 in Extended Data. For brain-behavior analysis regression tables, see Tables 3-4 and 3-5 in Extended Data.

To assess functional interactions between the dlPFC and mesolimbic system, we performed a psychophysiological interaction analysis (PPI) using the dlPFC as the seed and examined functional connectivity for high-relative to low-reward stimuli. We were specifically interested in examining dlPFC-VTA functional connectivity given prior work implicating this circuit in driving motivated behaviors (Ballard et al., 2011). We extracted functional connectivity estimates from the VTA for each participant and examined the relationship between dlPFC-VTA functional connectivity and high-reward memory benefits as a function of age, controlling for the high-reward source image category, using multiple linear regression. Here, we also found a significant association between dlPFC-VTA functional connectivity and high-reward specific source memory benefits across all participants (**β** = 0.055, *t*(84) = 2.76, *p* = 0.0071). Additionally, we observed a dlPFC-VTA functional connectivity-by-age interaction (**β** = 0.047, *t*(84) = 2.09, *p* = 0.040) showing that the relation between dlPFC-VTA functional connectivity and high-reward specific source memory benefits became stronger with increasing age (Figure 4). There were no significant effects of age alone or high-reward source image category (*p*s > 0.90) and no significant associations between dlPFC-VTA functional connectivity and high-reward gist source memory benefits after 24 hours (see Extended Data Tables 4-1 & 4-2). In a whole-brain dlPFC-seeded PPI analysis, no regions showed significant task-related functional coupling with the dlPFC. These data suggest that interactions between top-down encoding mechanisms and reward-processing regions may increasingly drive high-reward associative memory benefits to a greater extent with increasing age, into young adulthood.

**Figure 4.**
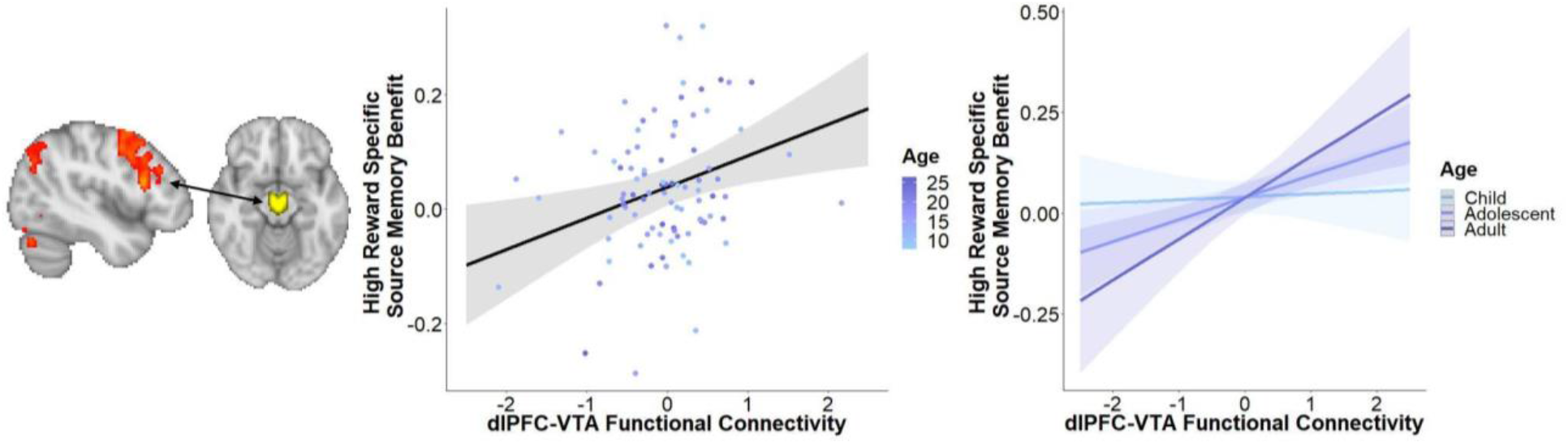
dlPFC encoding functional connectivity with VTA differentially relates to high-reward memory benefits after 24-hours across age. Increased dlPFC-VTA functional connectivity for high-relative to low-reward stimuli is associated with greater high-reward specific source memory benefits across all participants and this relationship becomes stronger with increasing age. Shading depicts a 95% confidence interval around predicted probability lines. Predicted probability lines are labeled with age groups for visualization purposes, but represent the mean and one standard deviation above and below the mean. The corresponding statistical analyses treat age as a continuous variable. For brain-behavior analysis regression tables, see Tables 4-1 and 4-2 in Extended Data.

### Experience-dependent changes in subcortical functional connectivity relate to high-reward memory benefits, particularly in younger individuals

We examined possible age-related changes in post-encoding consolidation mechanisms that might also facilitate the prioritization of high-reward associations in memory. We focused on experience-dependent changes in functional connectivity between the VTA and anterior hippocampus, which have been previously implicated in post-encoding reward memory processes (Gruber et al., 2016; Murty et al., 2017). We examined the relationship between pre- to post-encoding change in anterior hippocampus-VTA functional connectivity and high-reward memory benefits as a function of age using multiple linear regressions. There was no significant association between change in functional connectivity and high-reward specific source memory benefit after 24 hours (see Extended Data Table 5-1). We found a significant association between change in anterior hippocampus-VTA functional connectivity and high-reward gist source memory benefits (**β** = 0.17, *t*(81) = 2.01, *p* = 0.047). We also found a significant interaction between the change in functional connectivity and age (**β** = -0.20, *t*(81) = -2.35, *p* = 0.021), indicating that the relationship between change in anterior hippocampus-VTA functional connectivity and high-reward gist source memory benefits was stronger in younger participants (Figure 5). There was a marginal effect of high-reward source image category (**β** = -0.033, *t*(81) = -1.82, *p* = 0.073) and no significant effect of age alone (**β** = 0.0051, *t*(81) = 0.41, *p* = 0.68) (see Extended Data Table 5-2). These results suggest that increases in high-reward associative memory benefits are related to increased subcortical functional connectivity and that this association is largely driven by a stronger relationship in younger participants.

**Figure 5.**
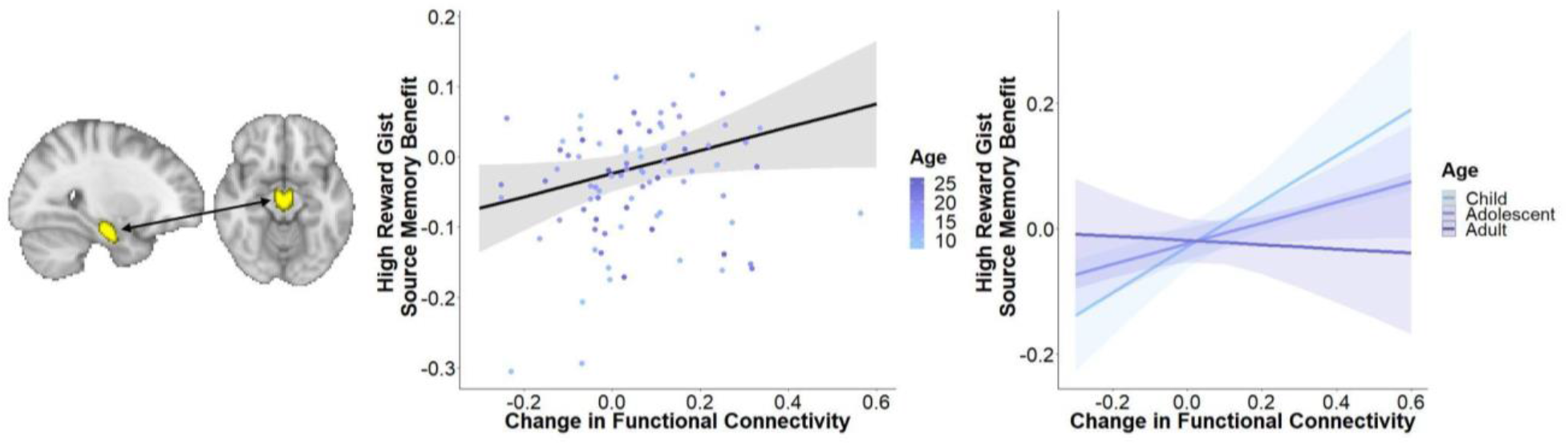
Post-encoding anterior hippocampus-VTA functional connectivity differentially relates to high-reward memory benefits after 24-hours across age. Increased change in anterior-VTA functional connectivity following reward-motivated encoding relative to baseline is associated with greater high-reward gist source memory benefits across all participants and this relationship is stronger in younger individuals. Shading depicts a 95% confidence interval around predicted probability lines. Predicted probability lines are labeled with age groups for visualization purposes, but represent the mean and one standard deviation above and below the mean. The corresponding statistical analyses treat age as a continuous variable. For brain-behavior analysis regression tables, see Tables 5-1 and 5-2 in Extended Data.

Taken together, we observed that encoding-related brain activation and functional connectivity were associated with high-reward specific source memory benefits, while post-encoding functional connectivity was associated with high-reward gist source memory benefits. One caveat is that while we see these distinct patterns of statistical significance with each memory measure, our analysis does not enable a direct statistical comparison between them, and thus we are not able to make strong claims about the selectivity of these brain-behavior relations. However, because specific and gist-level memory performance varied as a function of age but comprise a single associative memory process, our primary goal in this analysis was to identify age-related variation in the engagement of multiple neural processes known to collectively support reward-motivated associative memory formation.

## Discussion

The present study examined how reward influences the formation of long-term associative memories from childhood to adulthood, as the distributed neural circuits that support reward-motivated memory undergo dynamic changes. We found that after 24 hours, reward motivation enhanced specific source memories across participants of all ages and led to nonlinear age-related differences in general associative memory measures. Encoding-related functional connectivity between the VTA and dlPFC was associated with reward memory benefits, particularly in older participants. Post-encoding increases in functional connectivity between the VTA and anterior hippocampus were also associated with reward memory benefits, particularly in younger participants. Our findings point to possible developmental changes in the mechanisms through which reward motivation modulates associative memory and suggests that there may be age-varying contributions of post-encoding “offline” subcortical and “online” prefrontal cortical encoding processes supporting reward-motivated memory.

We found that reward-motivated associative memory enhancements consisted of both age-invariant and nonlinear age effects: while high-reward specific source memory was enhanced across all ages compared to low reward, we observed nonlinear age-related differences in general associative memory. High-reward general source memory was greatest during adolescence, however this effect seems to be driven by high-reward specific source memory performance because high- and low-reward gist source memory were similar during adolescence. Our results indicate that, in contrast to adolescents, younger and older individuals showed better general and gist source memory for low-reward associations. This pattern of age-related change mirrors a finding observed in a previous developmental study that examined a measure of general source memory for the outcomes of choices observed during a value-based learning task after 24-hours (see supplemental materials Katzman and Hartley, 2020). These converging findings may indicate that while children and adults show differential specificity and perhaps depth of processing (Craik, 2002) of encoding associations based on high- or low-reward level, adolescents may have specifically attended to and prioritized highly rewarded information in memory. This potential account is consistent with prior work that has demonstrated increased sensitivity to rewards during adolescence (Galván, 2013; Doremus-Fitzwater and Spear, 2016). Additionally, arousal states can modulate memory selectivity (see Clewett and Murty, 2019 for review), thus age-related differences in arousal responses to high- and low-reward stimuli during encoding may have contributed to the observed patterns of associative memory performance. Our findings suggest that although reward adaptively enhances memory for specific associations across all ages, developmental differences in reward sensitivity have unique consequences for more general associative memory.

Prefrontal encoding mechanisms also related to reward-motivated memory enhancements across age. Consistent with previous research in adults, we found increased brain activation in several cortical areas and the thalamus across participants of all ages for encoding high- relative to low-reward associations (Wittmann et al., 2005; Cohen et al., 2014; Murty and Adcock, 2014). We focused specifically on observed dlPFC activation given both prior work in adults underscoring its role in driving reward-motivated behaviors (Ballard et al., 2011), as well as evidence of its protracted developmental trajectory (Mills et al., 2014) and increasing contributions to memory encoding with age (Menon et al., 2005; Ofen et al., 2007; Ghetti et al., 2010; Nussenbaum and Hartley, 2021). We found that dlPFC activation was related to high-reward specific source memory benefits 24 hours after encoding across all participants. This age-invariant effect is consistent with intracranial recording evidence demonstrating that lateral PFC activity during encoding predicts subsequent memory across age in individuals ages six to nineteen years-old (Johnson et al., 2018). We also found that greater functional coupling between the dlPFC and VTA was related to greater high-reward specific source memory benefits after 24 hours across all participants, and that this association was stronger with increasing age. Given prior work indicating that prefrontal inputs to the VTA can instantiate reward motivation in adults (Ballard et al., 2011), our findings suggest that online, top-down modulation of the VTA strengthens with age, into adulthood. Our results suggest that the dlPFC can contribute to reward-motivated associative memory formation across all ages and that prefrontal-mesolimbic system encoding interactions more strongly contribute to the formation of reward memory with increasing age.

We also found evidence for age-related differences in subcortical post-encoding consolidation mechanisms that support reward-motivated memory enhancements. Experience-dependent increases in anterior hippocampus functional connectivity with the VTA from pre- to post- reward-motivated encoding related to high-reward gist source memory benefits after 24 hours, particularly in younger individuals. While we are unable to make strong claims about the selectivity of this effect, the relationship between more general associative memory and post-encoding functional connectivity may reflect the proposed role for consolidation mechanisms in abstraction and generalization processes (McClelland et al., 1995; Tambini and Davachi, 2019). Our results suggest that offline post-encoding subcortical interactions involving the mesolimbic system may more strongly contribute to the formation of reward memory in younger individuals. This result aligns with prior work in children that has shown greater sleep-dependent learning and memory enhancements and increased gains in recall of prior learning in children, relative to adults, after sleep (Kurdziel et al., 2013, 2018; Wilhelm et al., 2013). Both these studies of sleep and our work examining post-encoding processes during awake rest suggest that offline consolidation processes may be particularly consequential for the formation of long-term memories in children, as more rapidly deployable online encoding mechanisms continue to develop.

Our results are consistent with theoretical models of brain development that posit earlier development of connections within subcortical circuitry relative to connections between cortical and subcortical circuitry (Casey et al., 2016, 2019). While these models were initially proposed as neurobiological accounts of developmental change in emotional reactivity and regulation, we show that hierarchical changes in brain development may also have behavioral consequences for motivated memory processes. We find a stronger relationship between post-encoding subcortical functional connectivity and reward-motivated memory benefits in younger individuals and a stronger relationship between encoding cortical-subcortical functional connectivity and reward-motivated memory benefits in older individuals. Although directionality cannot be inferred from functional connectivity analyses, our results are consistent with proposals that prefrontal cortical modulation of subcortical circuitry continues to develop into adulthood (Casey et al., 2019). Top-down, online encoding mechanisms may drive reward-motivated memory benefits more effectively as descending prefrontal cortical projections functionally mature. Thus, prioritization of information in memory in younger individuals may initially rely more heavily on subcortical circuitry and come to rely more on systems-level circuitry with increasing age.

We initially hypothesized that we would observe adolescent-specific effects in reward-motivated memory processes. Although we observed nonlinear age differences in reward-related general associative memory, we did not find corresponding adolescent-specific effects in brain activation that related to this behavioral pattern. Prior work in rodents suggests that adolescent VTA neurons show smaller responses during reward-motivated learning, relative to adults, and stronger responses to stimuli previously associated with rewards after extinction learning (Kim et al., 2016). This finding suggests that we may expect to see adolescent-specific differences in mesolimbic system activity at longer timescales than those used in the present study. Moreover, it may be the case that shifting contributions within the distributed neural circuits implicated reward-motivated memory may give rise to adolescent-specific effects through complex mechanisms that are not neatly captured by relating functional connectivity within simple brain circuits to behavior. Future work is needed to directly test this possibility. Additionally, our current task design did not temporally isolate specific and general components of associative memory. Thus, we were not able to clearly establish the selectivity of neural signals related to these measures. Future studies examining the neural mechanisms underlying reward memory at different levels of specificity across development are needed. More broadly, the selective prioritization of specific high-reward associations in memory during adolescence may have important implications for reward memory-guided behavior (Murty et al., 2016).

In the present study, we demonstrate that reward motivation enhances memory via overlapping cognitive and neural routes with age-varying contributions from childhood to adulthood. These findings provide a foundation for future work exploring how motivationally salient information shapes the memories that drive motivated behaviors across age. Insights from this emerging area of research have potential implications for optimizing learning experiences across age (Fandakova and Bunge, 2016) and deepening our understanding of how motivated memories may contribute to the emergence of mental illness and substance abuse (Pittenger, 2013) during childhood and adolescence. Uncovering neural and cognitive mechanisms that underlie learning from and remembering motivational inputs across age may ultimately inform strategies that can be leveraged to shape memories for experiences and support healthy development.

## Acknowledgments

This work was supported by a Klingenstein-Simons Fellowship in Neuroscience, a Jacobs Foundation Research Fellowship, the NYU Vulnerable Brain Project, a National Science Foundation CAREER Award Grant No. 1654393 and a National Institute of Mental Health grant R01MH126183 (to C.A.H.); a National Science Foundation SBE Postdoctoral Research Fellowship Grant No. 1714321 and a National Institute on Drug Abuse grant K01DA053438 (to A.O.C.). We thank Betsy Deza, Anastasia Filimontseva, Hannah Walker, and James Wyngaarden for help with data collection, David Clewett and Kate Nussenbaum for helpful feedback on this manuscript, and the staff of NYU’s Center for Brain Imaging, especially Pablo Velasco, for technical support.

## Extended Data

**Figure 1-1.**
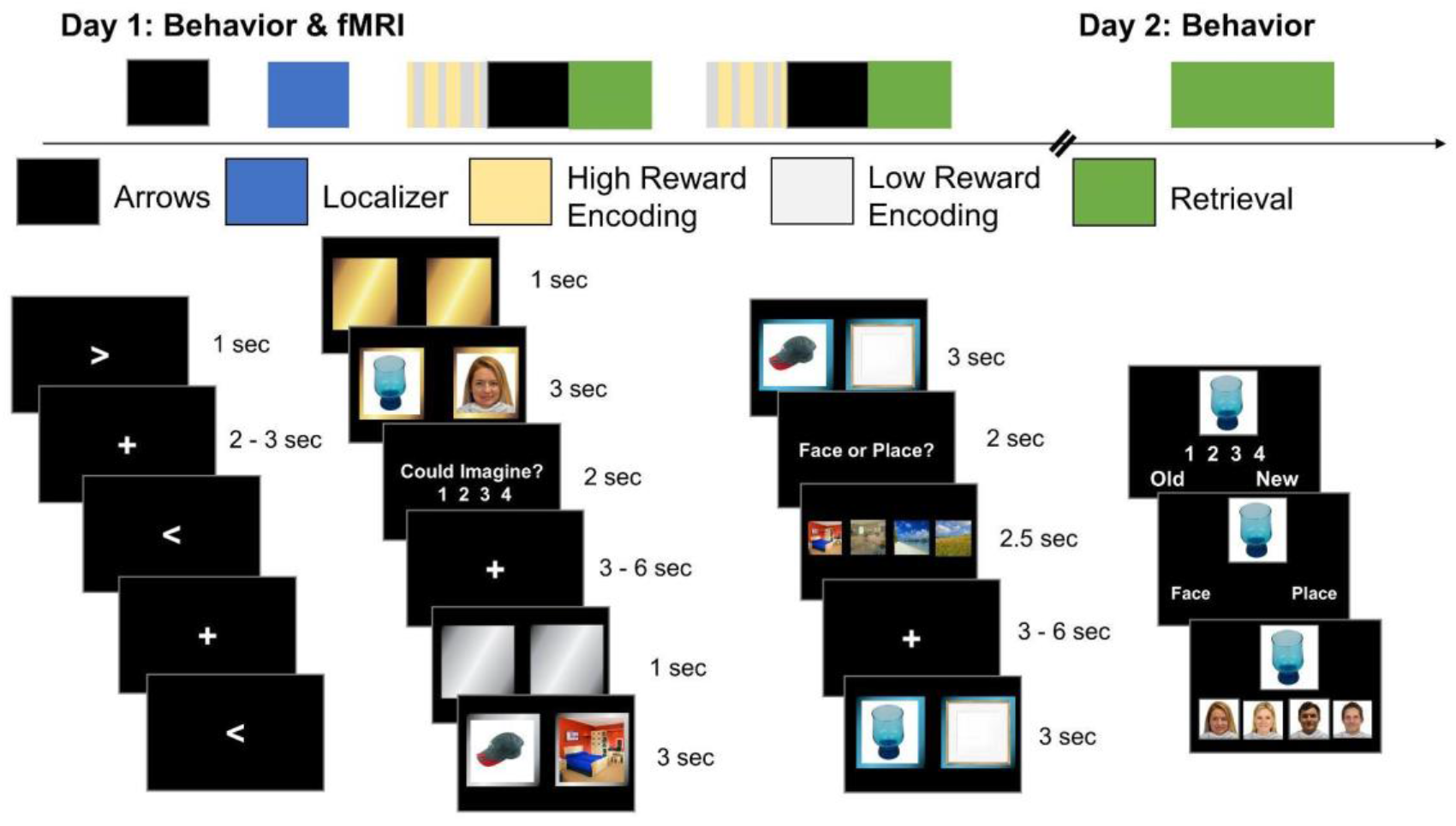
Task schematic including visualization of the scanned retrieval phase on day one. The scanned retrieval phase was included to address a different set of questions than those addressed in the present study.

**Figure 2-1.**
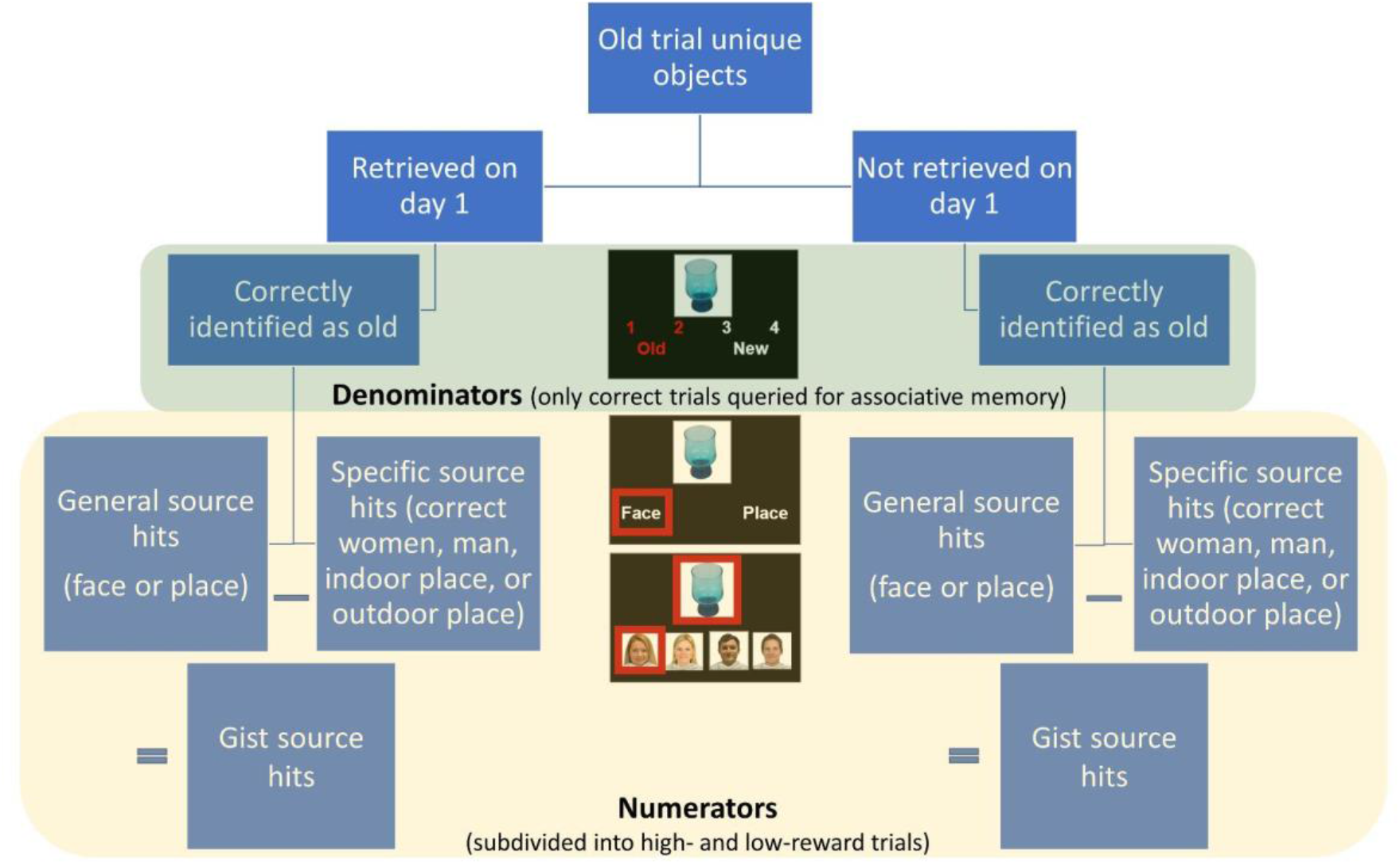
Visualization of how associative memory measures were computed.

**Figure 2-2.**
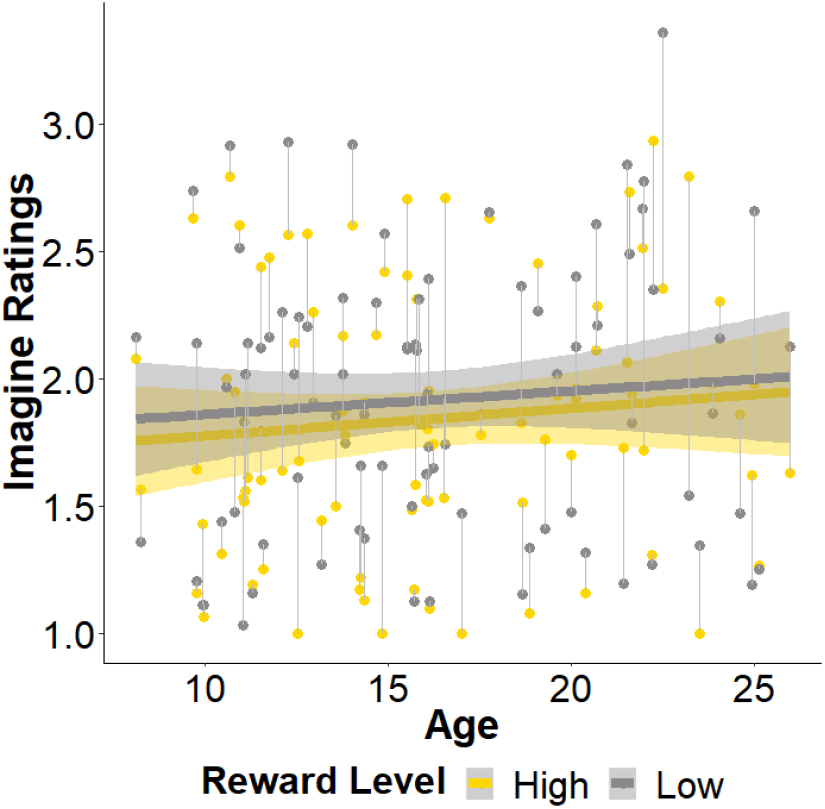
Imagine ratings during encoding. Using the same analysis approach as described for the associative memory measures, we conducted a linear mixed-effects analysis of ratings of how well participants imagined the story they told themselves about each pair of images. Ratings were on a scale from one (very easy to imagine) to four (very hard to imagine). While there was a trend towards a main effect of reward level on imagine ratings (*χ*2(1, N = 89) = 3.05, *p* = 0.081), such that individuals reported that they were better able to imagine high-reward associations across all ages, there were no significant effects of age (*χ*2(1, N = 89) = 0.79, *p* = 0.37), high-reward source image category (*χ*2(1, N = 89) = 0.81, *p* = 0.37), or reward-level by age interaction (*χ*2(1, N = 89) = 0.030, *p* = 0.86).

**Figure 2-3.**
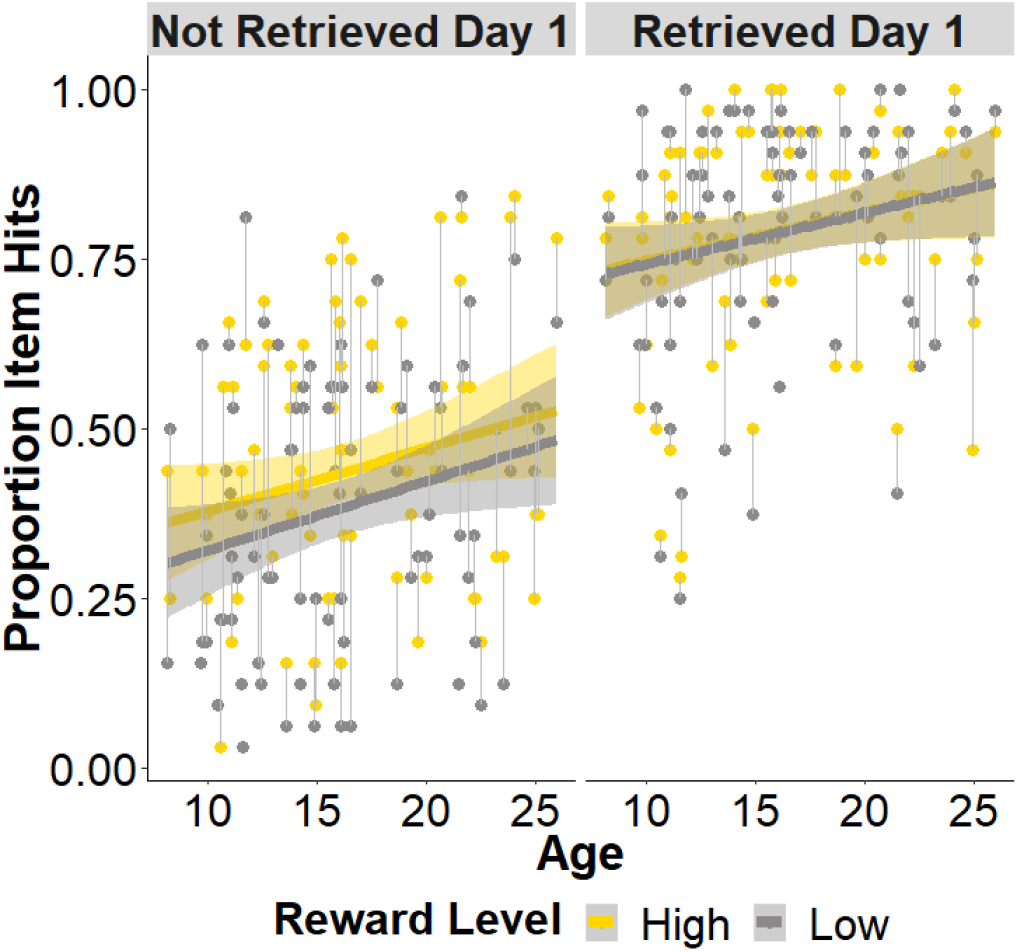
Item recognition memory after 24-hours. Using the same analysis approach as described for the associative memory measures, linear mixed-effects analysis of item recognition memory indicated significant main effects of reward level (*χ*2(1, N = 89) = 6.58, *p* = 0.010), retrieval condition (*χ*2(1, N = 89) = 1286.00, *p* < 0.0001), and age (*χ*2(1, N = 89) = 5.47, p = 0.020) that were qualified by a significant reward level-by-retrieval condition interaction (*χ*2(1, N = 89) = 5.44, *p* = 0.020). While we see a boost in memory for high-reward items across all ages for items that had not been retrieved on day one, we do not see significant differences between memory for high-and low-reward items that were retrieved on day one. This may be due to the fact that memory is at or near ceiling for a number of individuals. None of the associative memory measures show a reward level-by-retrieval condition interaction.

**Figure 2-4.**
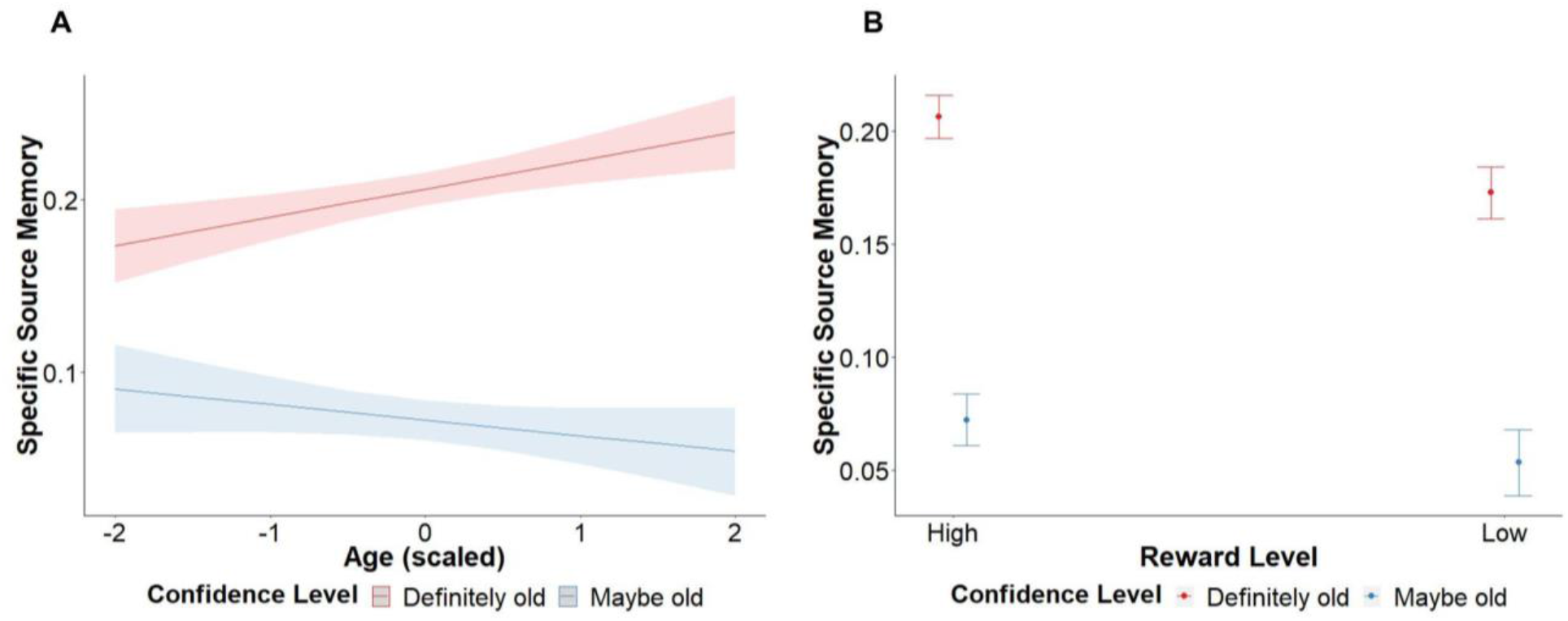
Item recognition memory confidence effects on specific source memory. (A) We observed a statistically significant confidence level-by-age interaction in specific source memory for paired associates involving items rated as “definitely old.” Specific source memory improved with increasing age, while specific source memory performance for paired associates involving items rated as “maybe old” was higher in younger than older individuals (*χ*2(1, N = 89) = 6.84, *p* = 0.0089). (B) There was also a statistically significant reward level-by-confidence interaction, such that the difference in memory performance between paired associates that involved items that were rated as “definitely old” and those that involved items that were rated as “maybe old” was greater for high-reward paired associates (*χ*2(1, N = 89) = 4.02, *p* = 0.045). Plots depict predicted values.

**Figure 2-5.**
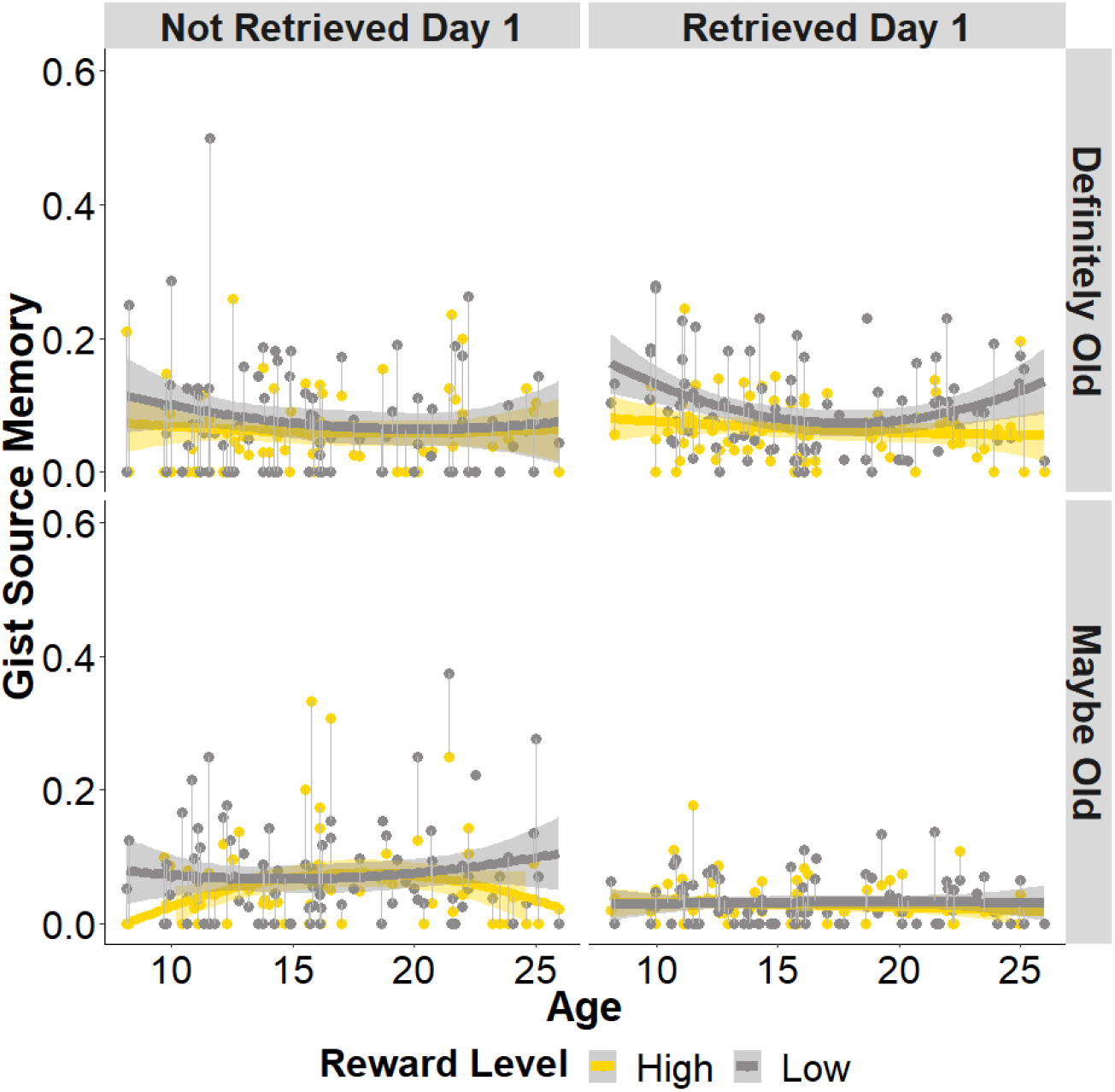
Item recognition memory confidence effects on gist source memory. We observed a statistically significant reward level-by retrieval condition-by-confidence level-by-age-squared interaction effect (*χ*2(1, N = 89) = 4.50, p = 0.034). This four-way interaction effect seems to be driven by more robust effects observed for paired associates that involved items that were rated as “definitely old” that were retrieved on day one and floor effects of paired associated that involved items that were rated as “maybe old” that were retrieved on day one, the latter likely due to a relatively small number of trials being identified as “maybe old.”

**Table 2-1.** General source memory analysis including fixed-effect of confidence level (definitely old or maybe old)

| *Predictors* | *Chisq* | *Df* | *P-value* |
| --- | --- | --- | --- |
| (Intercept) | 1272.024 | 1 | 2.2e-16 |
| Reward Level | 2.8276 | 1 | 0.09266 |
| Retrieval Condition | 2.1328 | 1 | 0.14418 |
| Age | 1.853 | 1 | 0.17344 |
| Confidence Level | 336.8276 | 1 | 2.2e-16 |
| Age^2 | 0.4376 | 1 | 0.50829 |
| High Reward Category | 0.0702 | 1 | 0.79101 |
| Reward Level*Retrieval Condition | 0.6231 | 1 | 0.42989 |
| Reward Level*Age | 0.6912 | 1 | 0.40576 |
| Retrieval Condition*Age | 0.2403 | 1 | 0.62399 |
| Reward Level*Confidence Level | 0.3794 | 1 | 0.53791 |
| Retrieval Condition*Confidence Level | 48.6488 | 1 | 3.062e-12 |
| Age*Confidence Level | 0.7234 | 1 | 0.39503 |
| Reward Level*Age^2 | 3.9373 | 1 | 0.04723 |
| Retrieval Condition*Age^2 | 0.3987 | 1 | 0.52777 |
| Confidence Level*Age^2 | 0.0064 | 1 | 0.93606 |
| Reward Level*Retrieval Condition*Age | 0.0022 | 1 | 0.96245 |
| Reward Level*Retrieval Condition*Confidence Level | 0.1428 | 1 | 0.70552 |
| Reward Level*Age*Confidence Level | 1.4253 | 1 | 0.23253 |
| Retrieval Condition*Age*Confidence Level | 0.4969 | 1 | 0.48085 |
| Reward Level*Retrieval Condition*Age^2 | 0.0815 | 1 | 0.77531 |
| Reward Level*Confidence Level*Age^2 | 0.0014 | 1 | 0.97031 |
| Retrieval Condition*Confidence Level*Age^2 | 0.0271 | 1 | 0.86925 |
| Reward Level*Retrieval Condition*Age*Confidence Level | 0.082 | 1 | 0.77465 |
| Reward Level*Retrieval Condition*Confidence Level*Age^2 | 0.5383 | 1 | 0.46313 |

**Table 2-2.** Specific source memory analysis including fixed-effect of confidence level (definitely old or maybe old)

| *Predictors* | *Chisq* | *Df* | *P-value* |
| --- | --- | --- | --- |
| (Intercept) | 822.9096 | 1 | 2.2e-16 |
| Reward Level | 9.4027 | 1 | 0.002167 |
| Retrieval Condition | 26.7884 | 1 | 2.27e-07 |
| Age | 1.9838 | 1 | 0.158989 |
| Confidence Level | 785.1638 | 1 | 2.2e-16 |
| High Reward Category | 0.7035 | 1 | 0.401622 |
| Reward Level*Retrieval Condition | 0.9715 | 1 | 0.324319 |
| Reward Level*Age | 0 | 1 | 0.99967 |
| Retrieval Condition*Age | 0.1076 | 1 | 0.742919 |
| Reward Level*Confidence Level | 4.016 | 1 | 0.045053 |
| Retrieval Condition*Confidence Level | 71.0157 | 1 | 2.2e-16 |
| Age*Confidence Level | 6.8419 | 1 | 0.008904 |
| Reward Level*Retrieval Condition*Age | 0.4029 | 1 | 0.525595 |
| Reward Level*Retrieval Condition*Confidence Level | 0.7531 | 1 | 0.385506 |
| Reward Level*Age*Confidence Level | 0.9968 | 1 | 0.31809 |
| Retrieval Condition*Age*Confidence Level | 0.0696 | 1 | 0.791896 |
| Reward Level*Retrieval Condition*Age*Confidence Level | 0.3498 | 1 | 0.55422 |

**Table 2-3.** Gist source memory analysis including fixed-effect of confidence level (definitely old or maybe old)

| *Predictors* | *Chisq* | *Df* | *P-value* |
| --- | --- | --- | --- |
| (Intercept) | 205.1212 | 1 | 2.2e-16 |
| Reward Level | 0.658 | 1 | 0.417256 |
| Retrieval Condition | 7.5022 | 1 | 0.0061623 |
| Age | 0.9929 | 1 | 0.3190289 |
| Confidence Level | 7.3245 | 1 | 0.006802 |
| Age^2 | 0.8312 | 1 | 0.361937 |
| High Reward Category | 1.317 | 1 | 0.251124 |
| Reward Level*Retrieval Condition | 0.1505 | 1 | 0.6980839 |
| Reward Level*Age | 1.4422 | 1 | 0.2297809 |
| Retrieval Condition*Age | 1.8233 | 1 | 0.1769175 |
| Reward Level*Confidence Level | 1.0008 | 1 | 0.317129 |
| Retrieval Condition*Confidence Level | 13.4504 | 1 | 0.0002449 |
| Age*Confidence Level | 8.6271 | 1 | 0.003312 |
| Reward Level*Age^2 | 7.1685 | 1 | 0.0074194 |
| Retrieval Condition*Age^2 | 1.2957 | 1 | 0.2549952 |
| Confidence Level*Age^2 | 6.1748 | 1 | 0.0129581 |
| Reward Level*Retrieval Condition*Age | 0.6391 | 1 | 0.4240536 |
| Reward Level*Retrieval Condition*Confidence Level | 0.1192 | 1 | 0.7298729 |
| Reward Level*Age*Confidence Level | 0.572 | 1 | 0.449449 |
| Retrieval Condition*Age*Confidence Level | 0.5898 | 1 | 0.4424863 |
| Reward Level*Retrieval Condition*Age^2 | 0.2318 | 1 | 0.6302254 |
| Reward Level*Confidence Level*Age^2 | 3e-04 | 1 | 0.9865002 |
| Retrieval Condition*Confidence Level*Age^2 | 0.0209 | 1 | 0.8851413 |
| Reward Level*Retrieval Condition*Age*Confidence Level | 0.3306 | 1 | 0.5652884 |
| Reward Level*Retrieval Condition*Confidence Level*Age^2 | 4.4969 | 1 | 0.0339561 |

**Figure 3-1.**
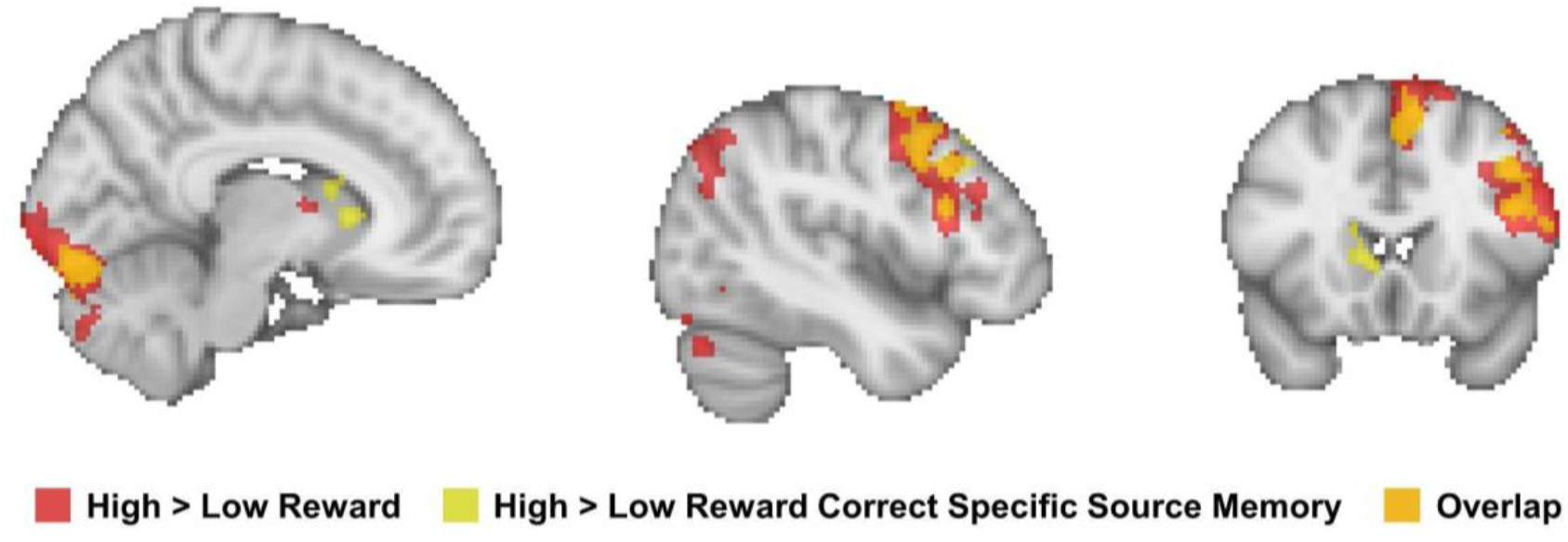
Encoding brain activation maps for high > low reward contrast, high > low reward restricted to trials with correct specific source memory after 24 hours, and their overlap. To assess the extent to which the reported encoding activations were reflective of successful memory encoding processes, the encoding fMRI analysis as described in the Materials & Methods section was rerun with the following four task regressors that reflect correct or incorrect specific souce memory after 24 hours: high-reward correct trials, low-reward correct trials, high-reward incorrect trials, and low-reward incorrect trials. Due to the presence of an empty correct trial task regressor in a run, five participants lost runs and one of those participants (who already had partial data due to a run eliminated for excessive motion) had to be excluded from this follow-up analysis (thus *n* = 88). We assessed high > low reward correct as the contrast of interest. We then assessed the degree of overlap between the high > low reward contrast map (that included all trials irrespective of subsequent accuracy) and the high > low reward correct contrast map.

**Table 3-1.**
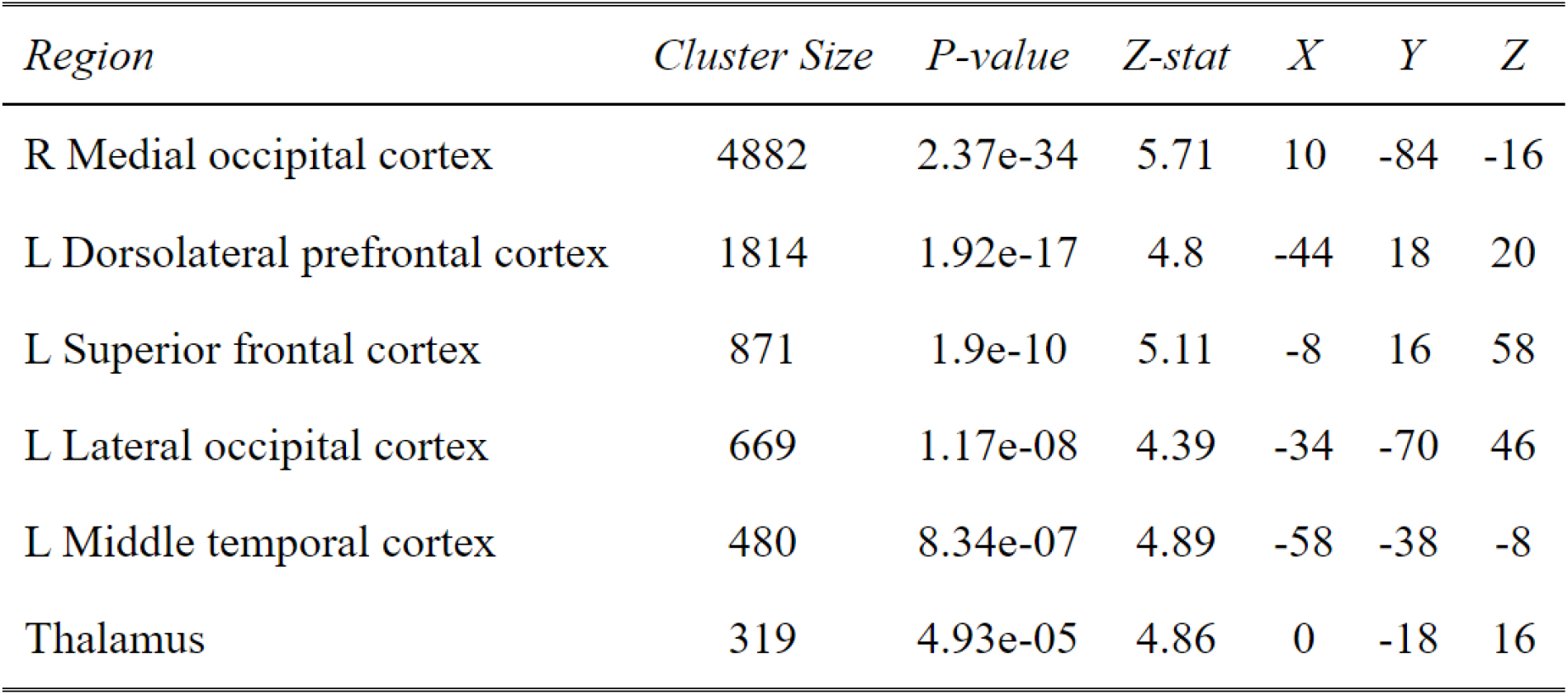
High > low reward activation map (p < 0.05, whole-brain corrected)

| *Region* | *Cluster Size* | *P-value* | *Z-stat* | *X* | *Y* | *Z* |
| --- | --- | --- | --- | --- | --- | --- |
| R Medial occipital cortex | 4882 | 2.37e-34 | 5.71 | 10 | -84 | -16 |
| L Dorsolateral prefrontal cortex | 1814 | 1.92e-17 | 4.8 | -44 | 18 | 20 |
| L Superior frontal cortex | 871 | 1.9e-10 | 5.11 | -8 | 16 | 58 |
| L Lateral occipital cortex | 669 | 1.17e-08 | 4.39 | -34 | -70 | 46 |
| L Middle temporal cortex | 480 | 8.34e-07 | 4.89 | -58 | -38 | -8 |
| Thalamus | 319 | 4.93e-05 | 4.86 | 0 | -18 | 16 |

**Table 3-2.** High > low reward correct specific source memory activation map (*p* < 0.05, whole-brain corrected)

| *Region* | *Cluster Size* | *P-value* | *Z-stat* | *X* | *Y* | *Z* |
| --- | --- | --- | --- | --- | --- | --- |
| R Medial occipital cortex | 853 | 2.19e-10 | 4.84 | 8 | -84 | -16 |
| L Dorsolateral prefrontal cortex | 617 | 5.96e-08 | 4.23 | -42 | 18 | 38 |
| L Superior frontal cortex | 453 | 1.37e-06 | 5.28 | -8 | 16 | 56 |
| R Frontal pole | 380 | 8.7e-06 | 4.83 | 28 | 56 | 6 |
| L Frontal pole | 285 | 0.00012 | 4.21 | -26 | 54 | -4 |
| R Paracingulate cortex | 190 | 0.0021 | 4.17 | 2 | 42 | 22 |
| R Caudate | 144 | 0.01 | 4.33 | 14 | 16 | 16 |
| Thalamus | 130 | 0.017 | 4 | 2 | -6 | 8 |
| L Superior frontal cortex | 123 | 0.022 | 3.88 | -22 | 24 | 48 |

**Table 3-3.** High > low reward correct specific source memory age main effect activation map (*p* < 0.05, whole-brain corrected)

| *Region* | *Cluster Size* | *P-value* | *Z-stat* | *X* | *Y* | *Z* |
| --- | --- | --- | --- | --- | --- | --- |
| R Primary motor cortex | 310 | 5.7e-05 | 4.43 | 2 | -14 | 52 |
| L Planum temporale | 165 | 0.0048 | 4.31 | -46 | -36 | 16 |

**Table 3-4.** dlPFC-high reward specific source memory benefit regression table.

| *Predictors* | **High-reward specific source memory benefit** |  |  |
| --- | --- | --- | --- |
|  | *Estimates* | *Conf. Int (95%)* | *P-Value* |
| Intercept | 0.03 | -0.01 – 0.06 | 0.155 |
| dlPFC activation | 0.03 | 0.01 – 0.06 | **0.003** |
| Age | -0.01 | -0.03 – 0.02 | 0.659 |
| High reward category | -0.01 | -0.06 – 0.03 | 0.542 |
| dlPFC activation*Age | -0.00 | -0.02 – 0.02 | 0.868 |

**Table 3-5.**
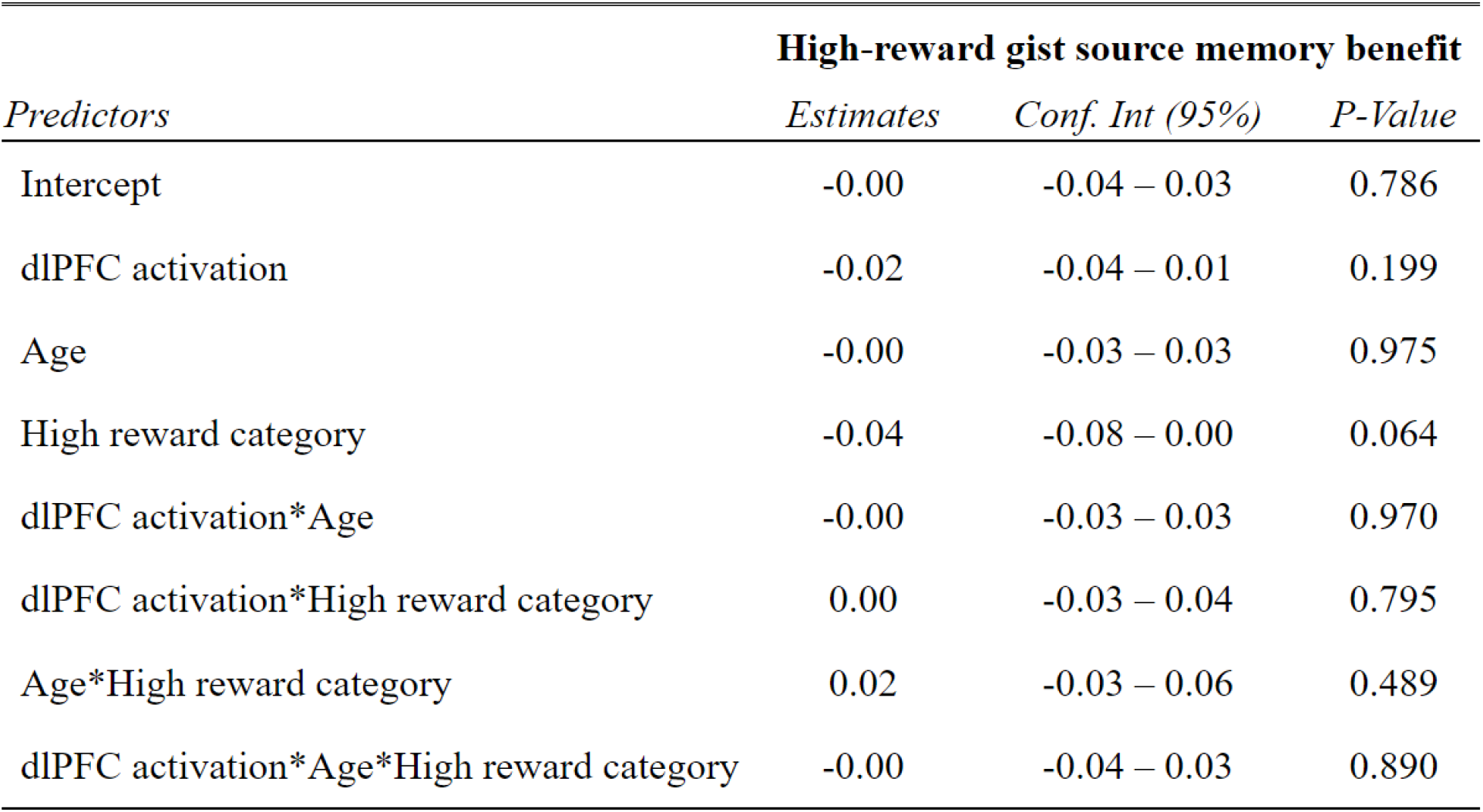
dlPFC-high reward gist source memory benefit regression table.

| *Predictors* | **High-reward gist source memory benefit** |  |  |
| --- | --- | --- | --- |
|  | *Estimates* | *Conf. Int (95%)* | *P-Value* |
| Intercept | -0.00 | -0.04 – 0.03 | 0.786 |
| dlPFC activation | -0.02 | -0.04 – 0.01 | 0.199 |
| Age | -0.00 | -0.03 – 0.03 | 0.975 |
| High reward category | -0.04 | -0.08 – 0.00 | 0.064 |
| dlPFC activation*Age | -0.00 | -0.03 – 0.03 | 0.970 |
| dlPFC activation*High reward category | 0.00 | -0.03 – 0.04 | 0.795 |
| Age*High reward category | 0.02 | -0.03 – 0.06 | 0.489 |
| dlPFC activation*Age*High reward category | -0.00 | -0.04 – 0.03 | 0.890 |

**Table 4-1.** dlPFC-VTA functional connectivity-high reward specific source memory benefit regression table.

| *Predictors* | **High-reward specific source memory benefit** |  |  |
| --- | --- | --- | --- |
|  | *Estimates* | *Conf. Int (95%)* | *P-Value* |
| Intercept | 0.04 | 0.01 – 0.07 | **0.021** |
| dlPFC-VTA connectivity | 0.05 | 0.02 – 0.09 | **0.007** |
| Age | -0.00 | -0.02 – 0.02 | 0.935 |
| High reward category | 0.00 | -0.05 – 0.05 | 0.996 |
| dlPFC-VTA connectivity*Age | 0.05 | 0.00 – 0.09 | **0.040** |

**Table 4-2.**
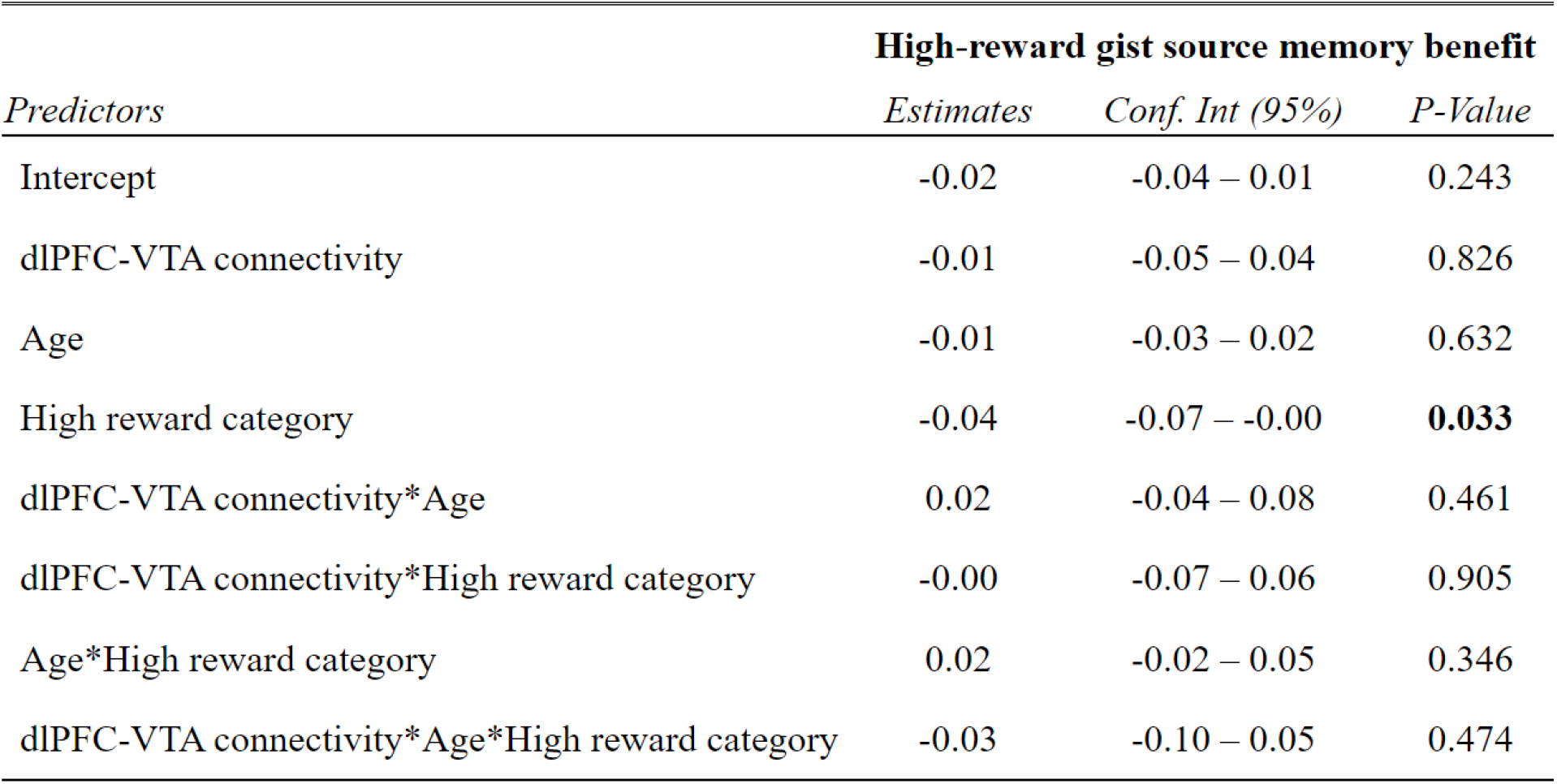
dlPFC-VTA functional connectivity-high reward gist source memory benefit regression table.

| *Predictors* | **High-reward gist source memory benefit** |  |  |
| --- | --- | --- | --- |
|  | *Estimates* | *Conf. Int (95%)* | *P-Value* |
| Intercept | -0.02 | -0.04 – 0.01 | 0.243 |
| dlPFC-VTA connectivity | -0.01 | -0.05 – 0.04 | 0.826 |
| Age | -0.01 | -0.03 – 0.02 | 0.632 |
| High reward category | -0.04 | -0.07 – -0.00 | **0.033** |
| dlPFC-VTA connectivity*Age | 0.02 | -0.04 – 0.08 | 0.461 |
| dlPFC-VTA connectivity*High reward category | -0.00 | -0.07 – 0.06 | 0.905 |
| Age*High reward category | 0.02 | -0.02 – 0.05 | 0.346 |
| dlPFC-VTA connectivity*Age*High reward category | -0.03 | -0.10 – 0.05 | 0.474 |

**Table 5-1.**
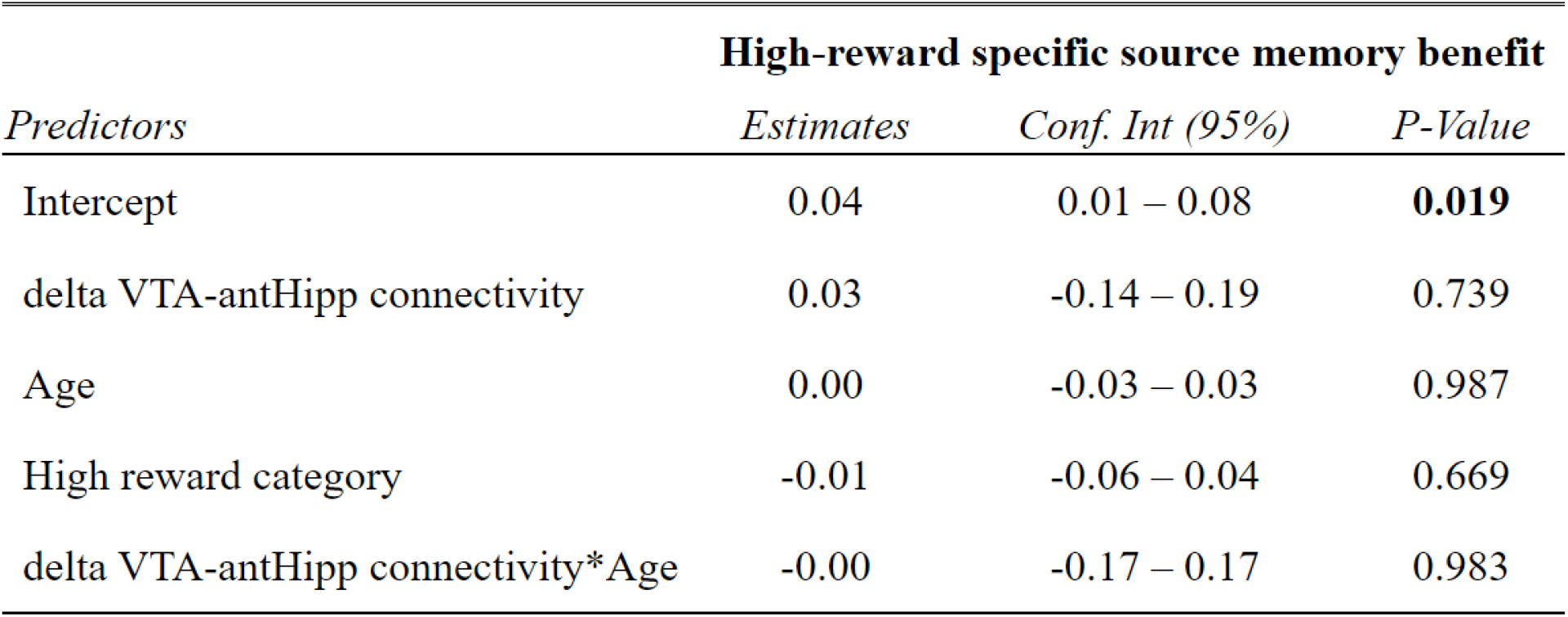
Change in VTA-anterior hippocampus functional connectivity-high reward specific source memory benefit regression table.

**Table 5-2.**
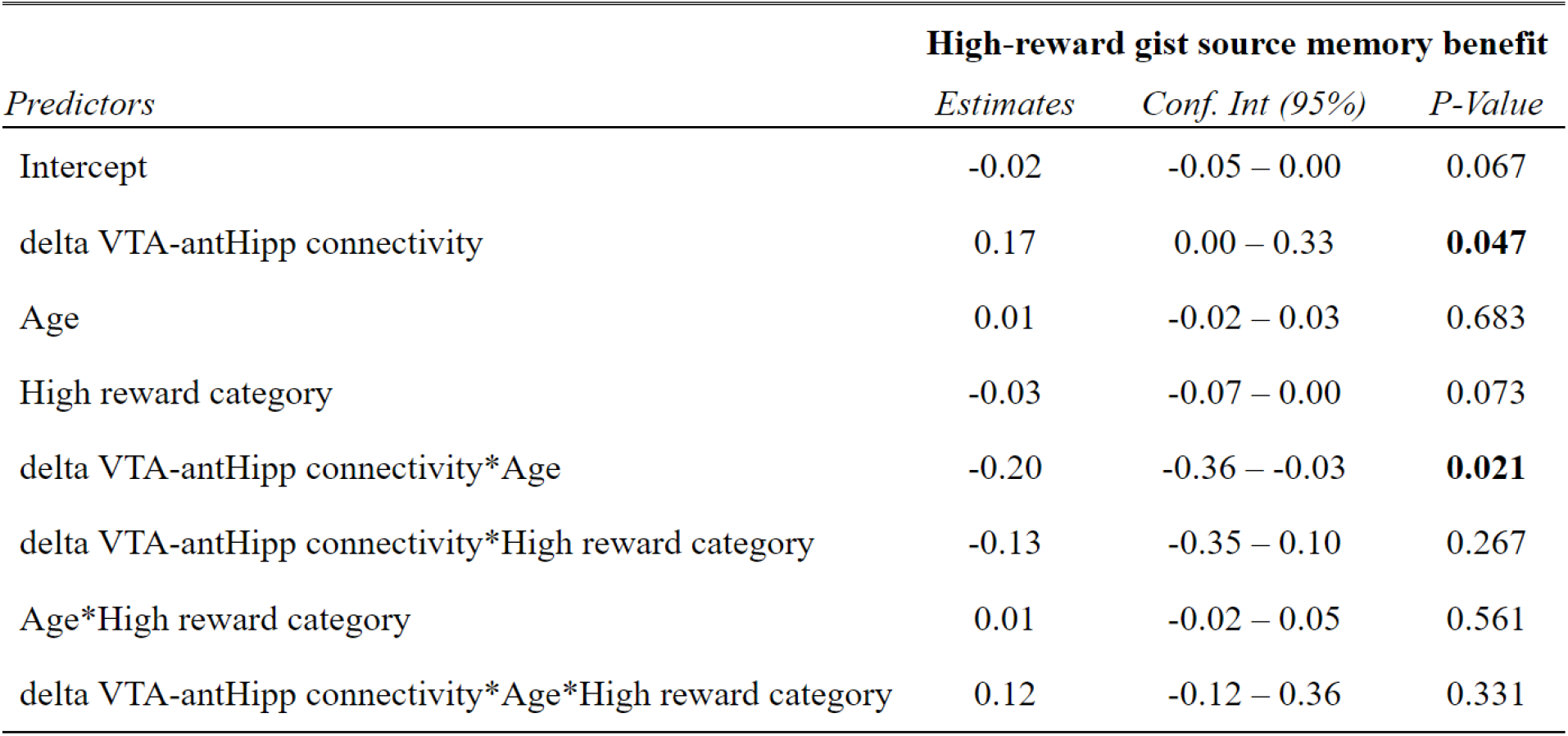
Change in VTA-anterior hippocampus functional connectivity-high reward gist source memory benefit regression table.

| *Predictors* | **High-reward gist source memory benefit** |  |  |
| --- | --- | --- | --- |
|  | *Estimates* | *Conf. Int (95%)* | *P-Value* |
| Intercept | -0.02 | -0.05 – 0.00 | 0.067 |
| delta VTA-antHipp connectivity | 0.17 | 0.00 – 0.33 | **0.047** |
| Age | 0.01 | -0.02 – 0.03 | 0.683 |
| High reward category | -0.03 | -0.07 – 0.00 | 0.073 |
| delta VTA-antHipp connectivity*Age | -0.20 | -0.36 – -0.03 | **0.021** |
| delta VTA-antHipp connectivity*High reward category | -0.13 | -0.35 – 0.10 | 0.267 |
| Age*High reward category | 0.01 | -0.02 – 0.05 | 0.561 |
| delta VTA-antHipp connectivity*Age*High reward category | 0.12 | -0.12 – 0.36 | 0.331 |

